# Genome-wide RNA structure changes during human neurogenesis drive gene regulatory networks

**DOI:** 10.1101/2021.08.02.454835

**Authors:** Jiaxu Wang, Tong Zhang, Zhang Yu, Wen Ting Tan, Ming Wen, Yang Shen, Finnlay R.P. Lambert, Roland G. Huber, Yue Wan

## Abstract

The distribution, dynamics and function of RNA structures in human development is under- explored. Here, we systematically assayed RNA structural dynamics and its relationship with gene expression, translation and decay during human neurogenesis. We observed that the human ESC transcriptome is globally more structurally accessible than that of differentiated cells; and undergo extensive RNA structure changes, particularly in the 3’UTR. Additionally, RNA structure changes during differentiation is associated with translation and decay. We also identified stage-specific regulation as RBP and miRNA binding, as well as splicing is associated with structure changes during early and late differentiation, respectively. Further, RBPs serve as a major factor in structure remodelling and co-regulates additional RBPs and miRNAs through structure. We demonstrated an example of this by showing that PUM2-induced structure changes on LIN28A enable miR-30 binding. This study deepens our understanding of the wide-spread and complex role of RNA-based gene regulation during human development.

## Introduction

Neuronal development is an extremely complex process that involves extensive gene regulation. A comprehensive understanding of how cells are regulated is needed to decipher the complexity of our brain. Beyond the primary sequence and RNA expression levels, recent developments have shown the importance of RNA structures in almost every step of the RNA lifecycle(Ganser et al., 2019), regulating cellular processes including transcription(Dethoff et al., 2012), translation(Kozak, 2005; Mao et al., 2014), localization(Martin and Ephrussi, 2009) and decay(Garneau et al., 2007). In addition to the static landscape of RNA structures inside cells, studying RNA structure dynamics, its regulators and cellular functions is key to understanding RNA structure function relationships. Recent high throughput structure probing have interrogated RNA structures across different vertebrate processes including during zebrafish embryogenesis (Beaudoin et al., 2018; Shi et al., 2020b), upon virus infections(Mizrahi et al., 2018) and across cellular compartments(Sun et al., 2019). While RNA structures have been shown to be associated with severe neurological diseases(Bernat and Disney, 2015; Kolb and Kissel, 2011) and important for diverse neuronal processes, including directing mRNA localization to the tips of axons in neurons(Jung et al., 2012), the full extent of RNA-structure based gene regulation during neuronal development is still largely unclear.

Here, to understand the role of RNA structure dynamics during human neuronal development, we utilized high throughput RNA structure mapping together with RNA sequencing, ribosome profiling and RNA decay studies for combinatorial analysis. We compared different analytic strategies to best identify structural changes between cellular states using RNA footprinting experiments as ground truth and showed that RNA structures are highly accessible in hESCs and are dynamic between cell types. We further identified additional regulators important for structure changes and demonstrated the importance and complexity of structure-based gene regulation during human neuronal development.

## Results

To study RNA structural dynamics during human neurogenesis, we differentiated hESCs into neurons using an established differentiation protocol (**Methods**). We performed high throughput RNA structure probing experiments *in vivo* using icSHAPE(Spitale et al., 2015), as well as other functional genomic experiments including RNA sequencing(Stark et al., 2019), ribosome profiling(Brar and Weissman, 2015) and RNA decay(Chen et al., 2008) in human embryonic stem cells (hESCs, Day 0), neural precursor cells (NPC, Day 7), immature neurons (iNeu, Day 8) and neurons (Neu, Day 14, **Figure 1a, Supp. Figure 1a, Supp. Table 1, Methods**). The combination of these high throughput experiments enabled us to study the impact of RNA structure on gene expression, translation and decay as the stem cells differentiate.

**Figure 1.**
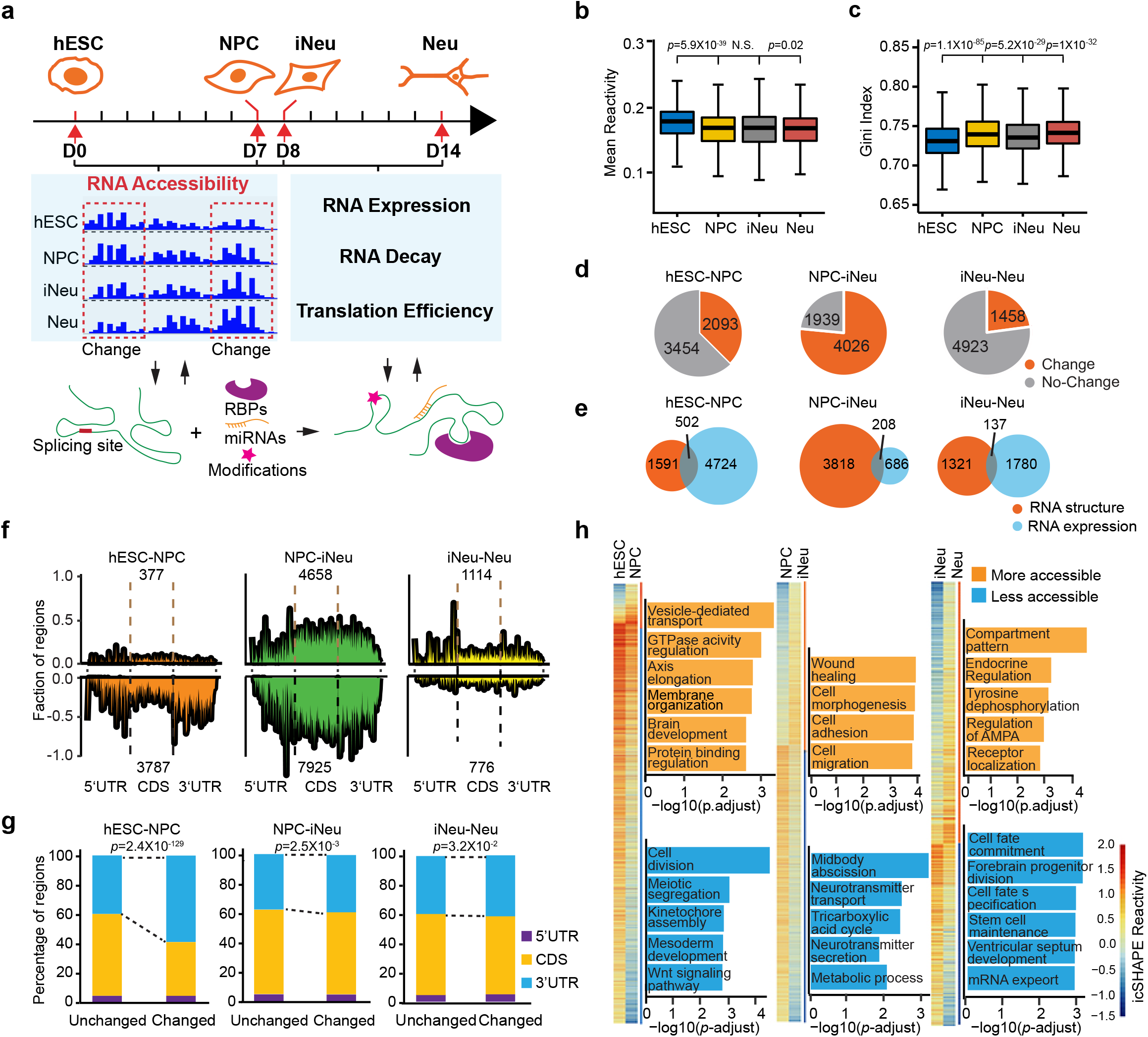
RNA structure profiling during neural development. **a,** Schematic of our experimental workflow. **b, c**, Boxplots showing the distribution of icSHAPE reactivity (**b**) and Gini index (**c**) per gene at the different time points of neuronal differentiation. The 3 comparisons were performed between neighboring time points. *P*-values were computed using two-sided Student’s T-test. **d,** Pie charts showing the numbers of genes with (orange) and without (grey) significant reactivity changes during neuronal differentiation. **e,** Venn diagram showing the overlap of RNA structure changed genes and RNA expression level changed genes during neurogenesis. **f,** The relative proportions of significantly changed regions on mRNAs that are becoming more accessible (upper panel) and less accessible (lower panel) between hESC-NPC (orange), NPC- iNeu (green), and iNeu-Neu (yellow). The number of changing regions is stated for both the upper and lower panels. The lengths of 5’ UTR, CDS and 3’ UTR are scaled to the same length. **g,** Stacked bar plots showing the numbers of windows without (left bar in each plot) and with (right bar in each plot) significant reactivity changes located on the 5’ UTR (purple), CDS (yellow) and 3’ UTR (blue). The *p*-values were computed using hypergeometric test. **h,** Left, heatmap showing the reactivity of structure changing windows at different stages. Right, GO enrichment of windows with significant reactivity changes between stages. Enriched GO terms in windows that are becoming more accessible are shown in yellow, while enriched GO terms in windows that are becoming less accessible are shown in blue.

We observed that RNA reactivity signals from NAI-N3- and DMSO-treated cells cluster independently from each other as expected (**Supp. Figure 1b**). icSHAPE reactivity, gene expression, translation and decay experiments showed high correlation between biological replicates (*R*>=0.97, 0.99, 0.89, 0.99 respectively) in each of the different time points, indicating that our high throughput measurements are reproducible (**Supp. Figure 1c-f**). Additionally, GO term analysis of transcripts that undergo TE (translation efficiency), decay and gene expression changes during neuronal differentiation showed that they are enriched for neuronal processes as expected (**Supp. Figure 1g**). We detected 3867, 4881, 5598, 5521 genes at hESC, NPC, iNeu and Neu stages using all four methods respectively (**Supp. Figure 2a, b, Supp. Table 2**). We observed a total of 2910 genes that are shared and successfully detected in all stages using the four methods (**Supp. Figure 2c, d**), indicating that we are interrogating a large fraction of the transcriptome.

Metagene analysis of icSHAPE reactivities on the hESC transcriptome showed that we could capture previously reported structural patterns in human transcriptomes(Kertesz et al., 2010), including the presence of a three nucleotide periodicity in the coding region and a negative correlation between Gini index and translation efficiency for the transcriptome (**Supp. Figure 3a, b**). Gini index is a measure of structural heterogeneity along a transcript, whereby a high Gini index indicates increased heterogeneity in icSHAPE reactivity and increased structure. Interestingly, our structure probing experiments showed that hESC RNAs have the highest icSHAPE reactivities and lowest Gini index as compared to RNAs from differentiated cells (**Figure 1b, c**), indicating that hESC RNAs are more structurally accessible. This result holds even when we calculate the icSHAPE reactivities for transcripts that are highly expressed across all four stages (**Supp. Figure 4a, b**). We also did not observe a good correlation between average reactivity and gene expression along a transcript (R=0.11), indicating that the increased reactivity of transcripts in hESC is not due to their increased transcript abundance (**Supp. Figure 4c**). Additionally, we validated this increase in RNA accessibility in hESC versus differentiated cell using two orthogonal structure probing methods, SHAPE-MaP and DMS-MaP-Seq (**Supp. Figure 4d-f**), confirming that hESC RNAs are indeed more structurally accessible than differentiated cells.

While the static picture of RNA structure landscapes has been mapped in many transcriptomes, being able to accurately detect structure changes remains to be a challenge. To find a statistically robust way to identify RNA structure changes sensitively and accurately across various time points, we compared different strategies of analyzing icSHAPE changes using RNAs that are known to change structure. This includes six riboswitch and two riboSNitch RNAs (**Supp. Figure 5a**) that are known to change structure in the presence of ligands or mutations respectively. We performed icSHAPE and traditional footprinting (the gold standard for in solution structure probing) to determine the analytic method that best captures the structure changes observed by footprinting (**Supp. Figure 5b, Supp. Table 3**). Both diff-BUMHMM and T-Test have been previously used to identify RNA structure changes in the transcriptome, however we observed that NOISeq outperforms both methods in its ability to identify structure changing regions sensitively and accurately (**Supp. Figure 5c, Supp. Table 3**). We further tested different window sizes and the extent of fold changes required to call a region as differentially changing, and observed that a window size of 20 bases and a fold change of 1.5 times is optimal for calling differential reactivities as these parameters enable sensitive detection of structure dynamics with low amount of false-positives (**Supp. Figure 5d, e**). We used NOI-seq with these parameters for all of our downstream analyses.

Upon neuronal differentiation, we observed that 2093 genes (4164 regions), 4026 genes (12538 regions) and 1458 genes (1890 regions) showed reactivity changes between hESC-NPC, NPC- iNeu and iNeu-Neu differentiation, respectively (**Figure 1d, Supp. Figure 6a**). Most of the transcripts had 1-2 regions of reactivity changes (**Supp. Figure 6b**), and the reactivity changes rarely exceeds 20 bases in each region (**Supp. Figure 6c, d**). Transcripts with structure changes show a poor overlap with transcripts that undergo gene expression changes during differentiation, indicating that RNA structure dynamics add an additional layer of information during differentiation (**Figure 1e**). Most of the RNA structural changes from hESCs to NPCs showed a decrease in reactivity (**Figure 1f**), agreeing with our observation that hESCs are highly structurally accessible (**Figure 1b, Supp. Figure 4**). These reactivity changing regions are enriched in the 3’UTRs (**Figure 1g**), indicating that the 3’UTRs either become more double-stranded or have additional cellular factors such as miRNA and RBPs bound to them during differentiation. Interestingly, we observed the largest number of RNA reactivity changes (12538 regions) as NPCs differentiate into immature neurons, when cells become fixed in their path towards neuronal lineage. Reactivity changing regions during NPC to iNeu differentiation are still enriched in the 3’UTRs, and become less accessible during differentiation (**Figure 1f, g**). As the cells differentiate from immature to more mature neurons, we observed fewer reactivity changes in the transcriptome and now an increased propensity for RNA regions to become more accessible.

To determine whether the reactivity changes along transcripts have functional consequences, we analyzed changes in translation, expression, and decay of these transcripts during neuronal differentiation. We observed that reactivity changes during hESC to NPC differentiation is associated with changes in translation (**Supp. Figure 7a**), agreeing with previous literature that transcripts with an increase in accessibility are more highly translated. We also performed GO term enrichments to determine the functions of the transcripts with reactivity changes. Structurally dynamic RNAs are enriched for GO terms including axon elongation, brain development, neurotransmitter processes and cell division (**Figure 1h**), and transcripts with large reactivity changes (two or more windows) are enriched for cell cycle, proliferation, and stem cell maintenance (**Supp. Figure 7b**). These results further indicate that RNA structures may play important roles during neurogenesis.

To identify RNA elements that show continuous reactivity changes during neuronal differentiation, we performed K means clustering to group the reactivity changes into five clusters (**Supp. Figure 8a**). We then correlated these clusters with gene regulatory processes including gene expression, translation and decay. Cluster 1 contains RNA regions that show a decrease in icSHAPE reactivities (**Figure 2a**), suggesting that they become more double-stranded over time. Similar to our above observations, regions in cluster 1 are significantly enriched in 3’UTRs (**Figure 2b**) and the transcripts that they belong to show an enrichment for faster RNA decay and decreased translation during differentiation (**Figure 2c, Supp. Figure 8b**). These transcripts with changes in decay and translation are enriched for GO terms including regulation of progenitor cell differentiation, mRNA destabilization and cell cycle phase transition (for faster decay) and cell-cell adhesion (for decreased translation, **Supp. Figure 8c**). Clusters 2 and 4 show a biphasic change in reactivity whereby the RNA regions either become more accessible-, and then less accessible (Cluster 2), or less accessible followed by being more accessible during differentiation (Cluster 4, **Figure 2a**). These two clusters are not enriched for translation or decay. Clusters 3 and 5 contain regions that show an increase in reactivities, indicating more single-strandedness over time (**Figure 2a**). These regions are enriched in the coding region (**Figure 2b**), show a positive correlation with translation and a negative correlation with decay (**Figure 2c**), and the transcripts that they belong to are enriched for GO terms associated with neurogenesis related pathways (**Supp. Figure 8c**). We did not see an enrichment with gene expression for any of the clusters (**Figure 2c, Supp. Figure 8d**), agreeing with our above observation that RNA structure dynamics are gene expression independent (**Figure 1e, Supp. Figure 4c**).

**Figure 2.**
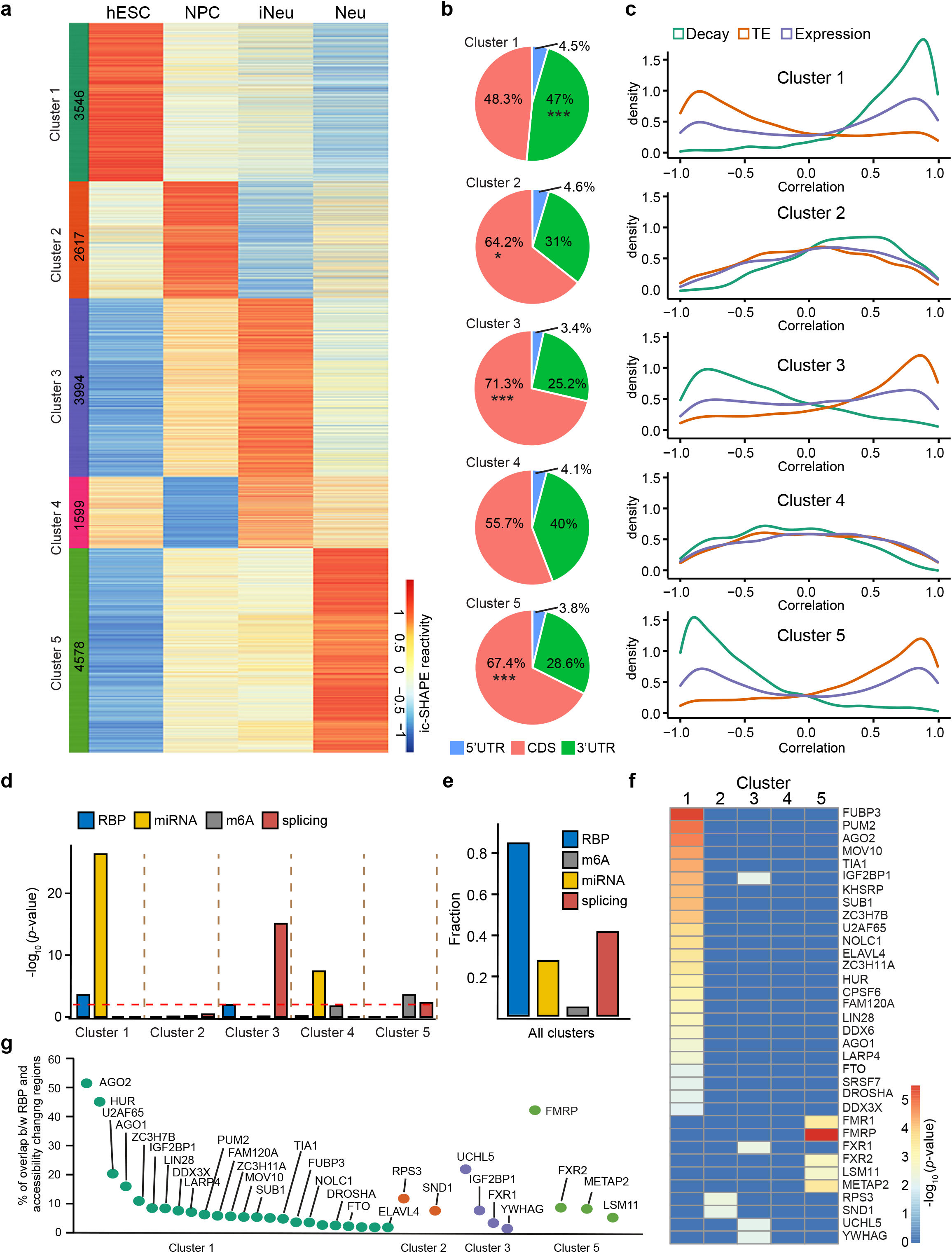
Functional analysis of structure clusters during neuronal differentiation. **a,** Heatmap showing K means clustering of reactivity changing windows across the four time points. The clustering was performed using Ward’s method based on the Euclidean distances on the normalized reactivity per window. **b,** Pie charts showing the proportion of windows localized in 5’ UTR (blue), CDS (red) and 3’ UTR (green) regions for each cluster. The enrichment test was performed using hypergeometric test. (*: *p*-value < 0.05; ***: *p*-value < 0.001). **c,** Density plots of Pearson’s correlation coefficient between normalized reactivity per windows and (1) RNA decay (green), (2) translation efficiency (orange) and (3) gene expression (purple) in each cluster. **d,** The enrichment of regulator binding sites including RBP binding sites (blue), miRNA binding sites (yellow), m6A sites (grey) and splicing sites (red) in each cluster. *P*-value is calculated using binomial test. The expected proportion is calculated as the overlapping proportion of different regulators to all windows in 5 clusters. **e,** The proportion of reactivity changing windows that overlap with RBP binding sites (blue), miRNA binding sites (yellow), m6A sites (grey) and splicing sites (red) in each cluster. Some of the reactivity changing windows overlap with two of more factors. **f,** Heatmap showing the –log10 of the enrichment *p*-values of RBP in each cluster. *P*-value was computed using hypergeometric test. **g,** The fraction of structure changing windows in each cluster that overlaps with the different enriched RBPs in that cluster.

Many cellular factors including miRNAs and RNA binding proteins regulate and are also in turn regulated by RNA structures(Kedde et al., 2010; Lewis et al., 2017; Liu et al., 2015; Spitale et al., 2015; Van Nostrand et al., 2020). To study the underlying mechanisms behind icSHAPE reactivity changes, we tested for the enrichment of miRNA, m6A modification, splicing and RBP binding sites near the reactivity changing windows in each cluster, as compared to all reactivity changing windows. Enrichments in the different clusters are independent of the window size used to overlap with the factors (**Supp. Figure 9a**). Cluster 1 is strongly enriched for miRNA and RNA binding protein sites (**Figure 2d, Supp. Figure 9a**), agreeing with our observation that it is associated with increased RNA decay (**Figure 2a-c**). Several of the enriched miRNAs, including miR-30 and miR-302, are important regulators of stem cell differentiation and stem cell pluripotency(Li et al., 2017; Rosa and Brivanlou, 2011) (**Supp. Figure 9b**). We also observed that cluster 3 is enriched for splice sites, indicating that splicing could be positively associated with increased translation during differentiation. Additionally, cluster 5, which shows a strong increase in reactivity over time (**Figure 2a**), is enriched for m6A modifications (**Figure 2d, Supp. Figure 9a**), again suggesting that m6A modifications can regulate translation during differentiation(Mao et al., 2019; Yu et al., 2018).

To determine the relative contributions of the different regulators in altering RNA structures, we calculated the proportion of reactivity changes that are attributed to each regulator. In the order of prevalence, 82%, 40%, 27% and 4% of the reactivity changing regions are associated with RBP, splicing, miRNA and m6A modifications respectively (**Figure 2e**), indicating that RBPs are a major contributor to reactivity changes. To identify the RBPs that are enriched in each cluster, we mined publicly available eCLIP datasets (**Methods**)(Van Nostrand et al., 2016a), and observed many enriched RBPs in different clusters (**Figure 2f**). Some of these RBPs could contribute to more than 10% of structural changes in a cluster (**Figure 2g),** suggesting that they play major roles in the remodeling of RNA structures. We found enrichment for RBPs that are functionally important for neuronal development, including PUM2, TIA1, IGF2BP1, ELAVL4, FTO, FMRP, FXR1, FXR2, Mov10 and FAM120A (**Figure 2f**). PUM2, TIA1 and ELAVL4, which were known to preferentially bind to 3’UTRs to regulate RNA stability are enriched in cluster 1(Ince-Dunn et al., 2012; Meyer et al., 2018; Wang et al., 2002), consistent with our observations that cluster 1 is positively correlated with RNA decay (**Figure 2c**). In addition, FMRP, which preferentially binds to the coding regions of RNA to regulate translation and decay in the literature(Darnell et al., 2011; Greenblatt and Spradling, 2018), is enriched in our cluster 5 (**Figure 2f, g**), which is associated with the coding region and with transcripts that show increased translation efficiency and decreased decay **(Figure 2c).** As some of these RBPs, including FMRP, Mov10 and FAM120A have been reported to be important in neurological diseases such as the Fragile X syndrome, suggesting that dysregulation of these RBP play roles in those disease through structure.

We also observed that cellular factors are enriched in different stages of the differentiation process, such that RBPs and miRNAs are enriched in reactivity changing regions during hESC to NPC differentiation, while splice sites are enriched as NPCs differentiate to neurons (**Figure 3a**). To comprehensively identify the RBPs that are associated with reactivity changing regions, we performed a more detailed search using sequence motif enrichments, and binding site enrichments using eCLIP datasets(Van Nostrand et al., 2016a) and binding sites predicted from the program PrismNet(Sun et al., 2021). We found a total of 35 enriched RBPs using eCLIP datasets, 68 RBPs using motif searches across different time points, and 51 using PrismNet (**Figure 3b, Supp. Figure 10a, Methods**). Overlap between eCLIP and motif sequences identified a total of ten shared RBPs, while four RBPs (PUM2, TIA1, CSTF2 and U2AF2) are consistently enriched across all three methods (**Supp. Figure 10b**). Nine out of ten RBPs enriched using eCLIP and motif sequences showed either an increase in gene expression or translation efficiency during hESC to NPC differentiation (**Supp. Figure 10c**), suggesting that an increase in their protein product could regulate the RNA structures of their targets.

**Figure 3.**
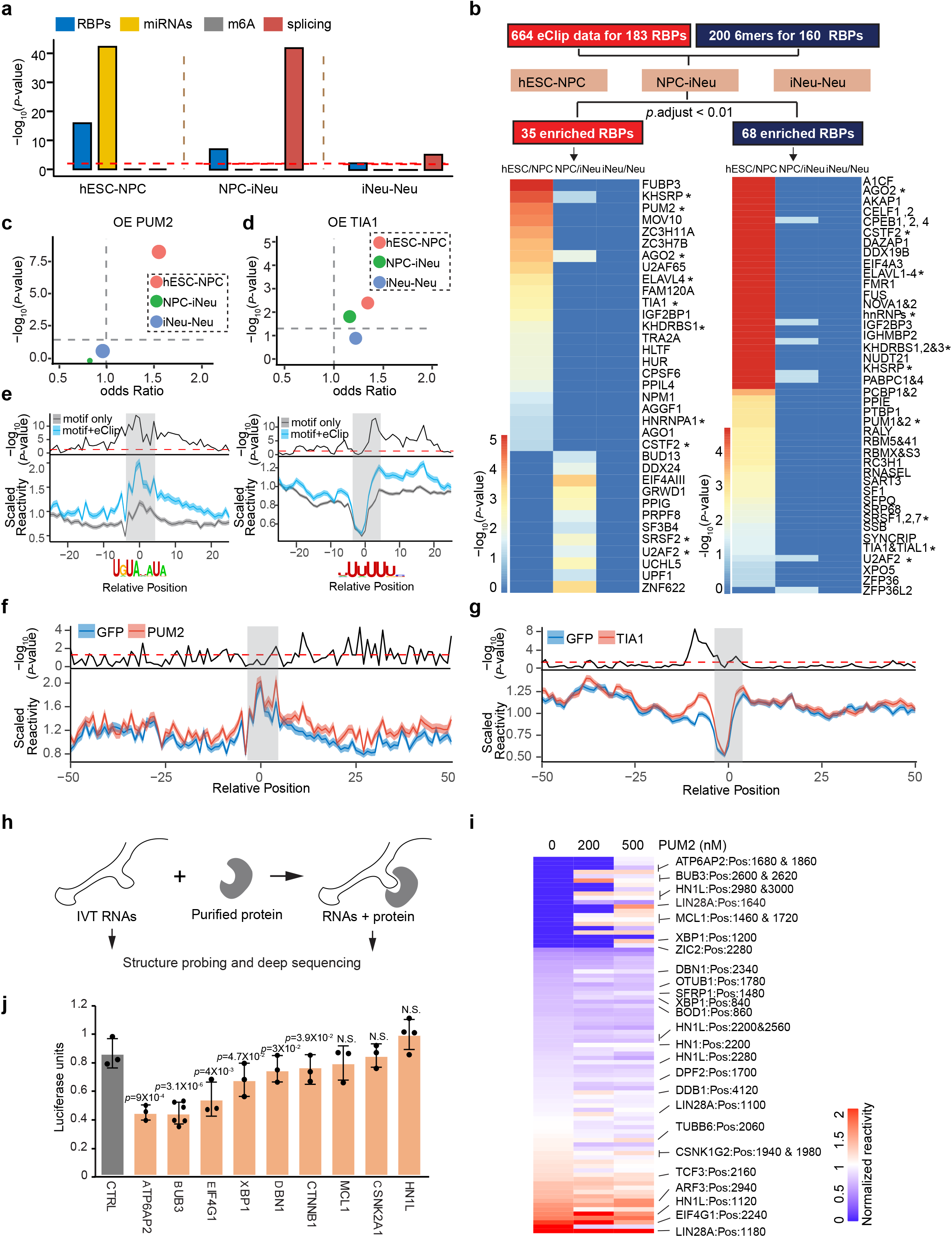
Reactivity changing regions are enriched for cellular factors. **a,** The enrichment of binding sites of cellular factors in reactivity changing windows during hESC-NPC, NPC-iNeu and iNeu-Neu differentiation. The number of reactivity changing regions that fall +/- 50 bases of the binding sites of RBPs (blue), miRNAs (yellow), m6A (grey) and splicing sites (red) are compared to all (changing and non-changing) windows that fall within +/- 50 bases of these factors. . *P*-value is calculated using binomial test. **b,** Heatmaps showing RBP enrichment in reactivity changing regions using eCLIP binding sites (left) and motif sequences (right). The 10 shared RBPs which are predicted by both methods are labeled with asterisk. **c, d,** Enrichment of significant reactivity changing windows during hESC-NPC, NPC-iNeu and iNeu-Neu with reactivity changing windows upon PUM2 (**c**) or TIA1 (**d**) overexpression. **e,** *Bottom*: Metagene analysis of normalized icSHAPE reactivity of real (with eCLIP and motif) and artificial (with motif only) PUM2 (left panel) and TIA1 (right panel) binding sites. *Top*: *P*-values on top of each subplots were calculated for every nucleotide using single-sided T-test. The grey shaded region indicates the location of the binding motif of PUM2(Hafner et al., 2010) and TIA1(Ray et al., 2013). Motif sequences are indicated below each subplot. **f, g,** Metagene analysis of normalized icSHAPE reactivity around real PUM2 (**f**) or TIA1 (**g**) binding sites before and after PUM2 (**f**) or TIA1 (**g**) overexpression. Real binding sites are sites with both motif sequences and eCLIP binding evidence. *P*-values on top of plot were calculated per nucleotide using the single-sided T-test. The grey shaded box indicates the location of the motif. **h,** Schematic of our *in vitro* RNA-protein interaction assay. **i,** Heatmap showing the normalized SHAPE-MaP reactivity of 87 RNA fragments incubated with different concentrations of PUM2 (0, 200, 500nM). **j,** Ratio of relative luciferase units of 9 randomly selected transcript regions cloned behind luciferase, upon PUM2 or GFP overexpression in HEK293T cells. *P*-values were calculated using the one-tail T-test, replicate number n=3∼6)

To understand the relationship between RBP and RNA structures, we focused our studies on two neurologically important RBPs- PUM2 and TIA1- which are enriched during hESC to NPC differentiation. Western blot experiments showed that the protein levels of both PUM2 and TIA1 are elevated upon differentiation (**Supp. Figure 10d**). To determine the impact of these two RBPs on transcript structures, we individually over-expressed them in hESCs using tetracycline-controlled transcription activation (Tet-on system, **Methods, Supp. Figure 11a-c**) and performed structure probing in RBP overexpressed and control hESC. Overexpression of PUM2 or TIA1 in hESC resulted in reactivity changes that significantly overlapped with naturally occurring reactivity changes during hESC to NPC differentiation (**Figure 3c, d**), indicating that PUM2 and TIA1 could regulate RNA reactivities during differentiation.

Studying the structural contexts of RBP targets enables us to understand the substrate specificity of RBPs(Dominguez et al., 2018). In addition, RBP binding can result in structure changes that have important functional consequences for their targets. While PUM2 is known to bind to single-stranded regions(Jarmoskaite et al., 2019), TIA1 is not known to have a clear structural context for its substrate. By comparing “real” binding sites (motifs with evidence of eCLIP binding) with “artificial” binding sites (motifs without evidence of eCLIP binding), we observed that real PUM2 motifs are more accessible around the binding motifs as compared to artificial motifs (**Figure 3e**). This agrees with previous literature that PUM2 prefers binding to more single-stranded regions(Jarmoskaite et al., 2019). Interestingly, real TIA1 binding sites are also more accessible than artificial sites and we also observed that TIA1 binding motifs contain lower reactivity regions as compared to its neighboring sequences (**Figure 3e**). This suggests that an accessible structural context of TIA1 motifs could facilitate its binding to its targets.

To determine the effect of protein binding on RNA structure, we performed metagene analysis of RNA reactivities at “real” PUM2 or TIA1 binding sites before and after their overexpression. Metagene analysis showed that PUM2 binding resulted in an increase in accessibility downstream of PUM2 motifs (**Figure 3f**), with 77% of PUM2 binding sites becoming more accessible upon PUM2 over-expression and 70% of sites becoming less accessible upon PUM2 knockdown (**Supp. Figure 11c-e**). Overexpression of TIA1 on the other hand resulted in a sharp, local increase in accessibility of 10 bases upstream of the “real” TIA1 motif (**Figure 3g**), with 65% of its targets becoming more accessible with increased TIA1 levels and 79% of its targets becoming less accessible with decreased TIA1 levels (**Supp. Figure 11a, b, d, f**). We confirmed that overexpression of PUM2 and TIA1 resulted in reactivity changes at their respective binding sites using predicted binding sites from the program PrismNet (**Supp. Figure 11g**).

To test the proportion of *in vivo* reactivity changes that are caused by direct RBP binding, we cloned 87 RNA regions from genes that showed different extents of structural changes upon PUM2 over-expression (**Supp. Figure 12**). We *in vitro* transcribed and refolded the RNAs as a pool and performed RNA-protein interaction experiments by incubating the RNAs with different concentrations of purified PUM2 protein *in vitro* (**Figure 3h, Methods**). 53% of our selected RNA regions showed an increase in reactivity upon PUM2 protein incubation *in vitro*, confirming our *in vivo* data that PUM2 protein binding indeed results in greater structural accessibility (**Figure 3f, i**). For nine of the regions that showed significant reactivity changes upon PUM2 binding in vitro, we additionally tested the effect of PUM2 binding on their gene expression inside cells. Using luciferase reporter assays, we observed gene repression in six out of nine RNA regions, indicating that PUM2 binding indeed alters the gene regulation of these RNA regions (**Figure 3j**).

While multiple RBPs can bind to an RNA to coordinate gene regulation, the extent to which their coordination is structure dependent is largely unknown. To determine whether RBP-induced reactivity changes could alter the accessibility of nearby regulators, we calculated the enrichment of RBP pairs near reactivity changing windows versus non-reactivity changing windows in our dataset (**Figure 4a, Methods, Supp. Table 4**). Enriched RBP pairs share similar transcript targets (**Supp. Figure 13a**), their structure changing regions are enriched in the 3’UTR of their targets (**Figure 4b**). We observed that enriched RBP pairs include PTBP1 and QKI (**Figure 4a**), which are master regulator of neuronal cell fates(Hardy et al., 1996; Shu et al., 2017; Xue et al., 2013), agreeing with their importance in our data.

**Figure 4.**
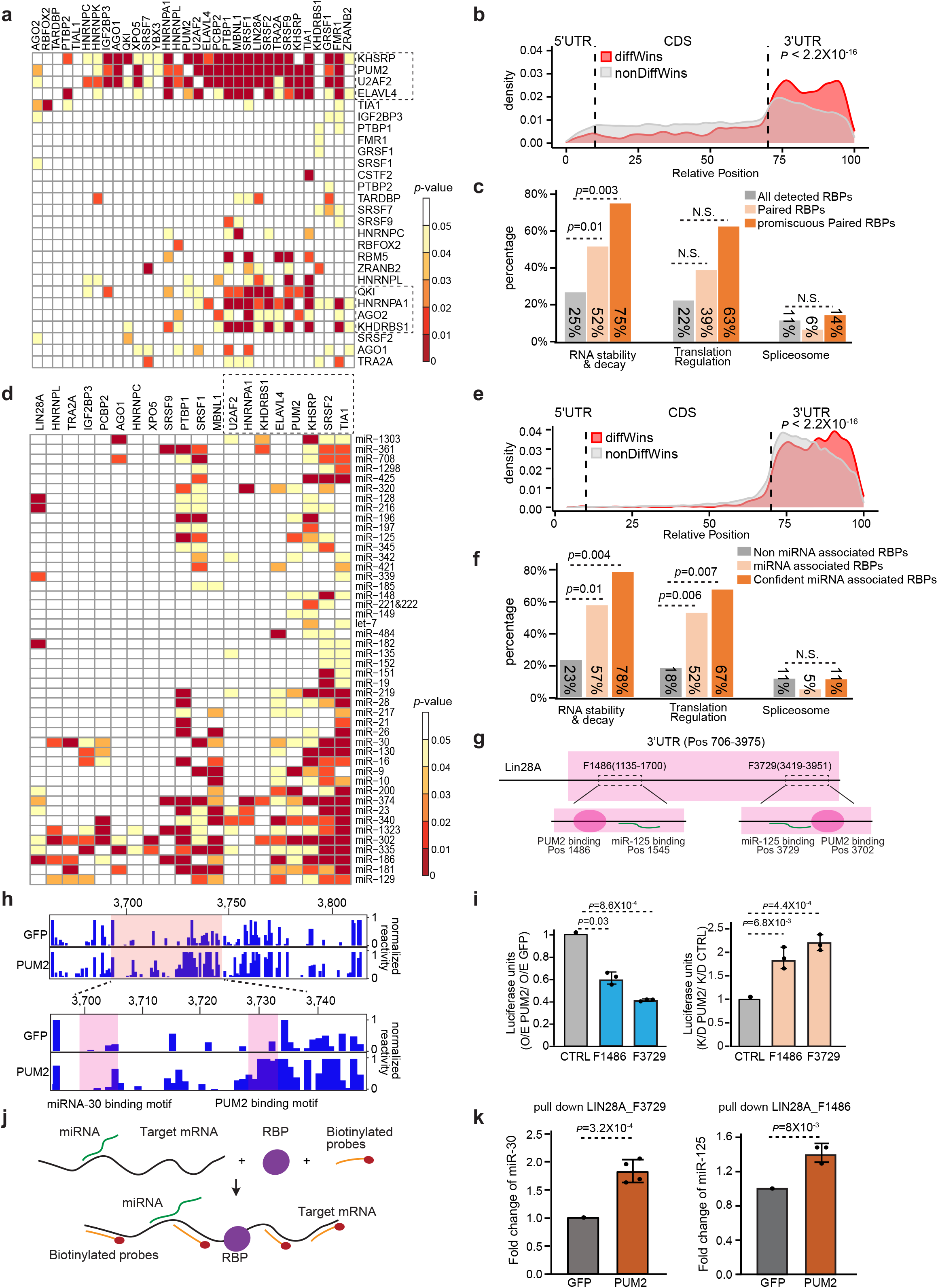
RBPs regulate the binding of other cellular factors through structure. **a,** Heatmap showing the enrichment of expressed secondary RBPs (shown as row names) in reactivity changing versus non-changing regions near a primary RBP binding sites (shown as column names) . *P*-values were calculated using hypergeometric test for RBPs that are enriched in reactivity changing regions +/- 100 bases around our enriched real RBP binding sites. The dotted boxes indicates the promiscuous RBPs (**Supp. Table 4**). **b,** Density plot of the location of structure changing windows (red) and the non-structure changing windows (grey) in significantly enriched RBP-RBP pairs along the transcriptome. The representative transcript is separated into 5’UTR, CDS and 3’UTR with a scaling ratio of 10:60:30. *P*-value was calculated using bootstrap Kolmogorov-Smirnov test. **c,** Barplots showing the function annotation of detected RBPs, paired RBPs and promiscuously paired RBPs. The functional annotation of individual RBPs is shown in Supp. Table 4. **d,** Heatmap showing the enrichment of top 100 expressed miRNAs in hESCs. 47 miRNAs with an enrichment to at least 2 RBPs were shown in heatmap. *P*-values were calculated using hypergeometric for miRNAs enriched in reactivity changing regions +/- 100 bases around enriched real RBP binding sites. The dotted box indicates the confident RBPs are enriched RBPs from both eCLIP and motif enrichment analysis from Figure 3b. **e,** Density plot of the location of structure changing windows (red) and the non-structure changing windows (grey) in significantly enriched RBP-miRNA pairs along the transcriptome. The representative transcript is separated into 5’UTR, CDS and 3’UTR with a scaling ratio of 10:60:30. *P*-value was calculated using bootstrap Kolmogorov-Smirnov test. **f,** Barplots showing the functional annotation of non-miRNA associated RBPs, miRNA associated RBPs and confident miRNA associated RBPs (**Supp. Table 4**). **g,** Schematic showing two binding sites of PUM2 (named F-1486 and F-3729 in pink circles) and two miRNA binding sites (miR-125 and miR-30 in green curves) on gene LIN28A. **h,** icSHAPE reactivity profile of PUM2 binding site on LIN28A (F-3729, in orange background) upon GFP overexpression (upper panel) and PUM2 overexpression (lower panel). The detailed reactivity profiles in the region from 3700 to 3740 are expanded below. **i,** Ratio of relative luciferase units of two LIN28A regions cloned behind luciferase, upon PUM2 versus GFP overexpression (left) and upon PUM2 knockdown versus control (right) in HEK293T cells, empty vector was used as control. *P*-values were calculated using the one-tail T-test, replicate number n=3. **j,** Schematic showing the experiment design of LIN28A pull down assay. **k,** Ratio of miR-30 (left) or miR-125 (right) amount pulled down with LIN28A under PUM2 and GFP overexpression. *P*-values were calculated using the one-tail T-test, replicate number n=4(left) and n=3 (right).

While some RBPs are enriched for other RBPs, we also identified RBPs that are promiscuously enriched for many other RBPs (**Figure 4a**). Enriched RBPs, and in particular promiscuously binding RBPs, are associated with functions such as regulation of RNA decay (**Figure 4c**). To determine whether target transcripts with RBP pairs located in reactivity changing regions are regulated differently from transcripts with RBP pairs found in non-reactivity changing regions, we determined changes in expression, decay and translation of these targets. We observed changes in target gene regulation when RBP pairs are associated with structure changes, highlighting that structure-based co-regulation contributes to an important layer of complexity in neuronal development (**Supp. Figure 13b**).

Besides RBPs, miRNAs also play important roles in gene regulation during neurogenesis. To determine whether RBP-induced reactivity changes could alter nearby miRNA binding as hESCs differentiate into NPCs, we analyzed the enrichment of top expressed miRNAs using sites predicted from TargetScan (**Methods, Supp. Table 4**). We observed 71 miRNAs that are enriched near RBP-associated reactivity changing regions (**Figure 4d**). These include miR-302, miR-30, miR-200, and miR-130, which are important miRNAs in stem cell maintenance, differentiation and iPS reprogramming(Li et al., 2017; Subramanyam et al., 2011; Wang et al., 2013). Enriched RBP-miRNA pairs have differential changes windows that are enriched in the 3’UTRs (**Figure 4e**) and are associated with functions including regulation of translation and decay (**Figure 4f**). These data suggest that RBP-induced structure changes could be a common mechanism for RBPs to regulate the binding of a second modulator to modulate gene expression.

To demonstrate the functional consequences of reactivity changes during neuronal differentiation, we focused our studies on RNAs that are functionally important and show large reactivity changes during the process. LIN28A, which is a key gene in stem cell maintenance, demonstrated the most number of structure changed regions upon PUM2 overexpression (**Supp. Figure 12**) and increased decay during differentiation(Shyh-Chang and Daley, 2013) (**Supp. Figure 14a, b)** which is consistent with previous report of Pum2 on the decay of its targets(Goldstrohm et al., 2018; Vessey et al., 2006; Wickens et al., 2002). We validated that PUM2 indeed binds to LIN28A in hESC and NPC cells using eCLIP (**Supp. Figure 14c**). PUM2 overexpression results in reactivity changes in more than thirty regions on LIN28A (**Supp. Figure 14d**), including an increase in icSHAPE reactivities around the PUM2 binding sites and two miRNA binding sites, miR-30 and miR-125, in the 3’UTR (**Figure 4g, h**). To confirm that these reactivity changes are indeed due to PUM2 binding to LIN28A, we incubated LIN28A with PUM2 *in vitro* and observed that the presence of PUM2 indeed increases RNA accessibility at PUM2, miR-30 and miR-125 binding sites (**Supp. Figure 14e, f**).

To determine the effect of PUM2 binding on LIN28A, we measured LIN28A levels in hESC and NPCs upon PUM2 knockdown. PUM2 knockdown resulted in an increase in LIN28A expression (**Supp. Figure 15a, b**), suggesting that PUM2 downregulates LIN28A. Additionally, we cloned each of the two LIN28A reactivity changing regions that are nearby PUM2 binding sites (**Figure 4g**), behind a luciferase reporter gene. Luciferase assays showed that overexpression of PUM2 decreased gene expression (**Figure 4i**), while downregulation of PUM2 increased gene expression of the two LIN28A regions, again indicating that LIN28A is downregulated in the presence of PUM2. As the PUM2-induced reactivity changes overlap with miR-30 and miR-125 binding sites, we hypothesized that reactivity changes during PUM2 over-expression or neuronal differentiation could impact the accessibility of these miRNAs to bind to LIN28A. Indeed, pull down of LIN28A using a pool of biotinylated antisense oligos upon PUM2 over-expression or during neuronal differentiated significantly increased the amount of miR-30 and miR-125 that bind to LIN28A (**Figure 4j, k, Supp. Figure 15c, d**).

To visualize the structure changes induced by PUM2 binding and its impact on the accessibility of miRNA binding on LIN28A, we focused on miR-30 as it was enriched in our data and is an important miRNA during stem cell differentiation. We incorporated icSHAPE reactivity as constraints in the Vienna RNAfold package to model LIN28A RNA secondary structures before and after PUM2 binding(Lorenz et al., 2011) (**Methods**). Interestingly, the structure models show extensive rearrangement, resulting in miR-30 binding site becoming more accessible, upon PUM2 binding (**Figure 5a**). To demonstrate that it is indeed PUM2-induced structure changes that impact miR-30 binding, we performed mutations on the LIN28A secondary structures to either destabilize the pre-PUM2 bound structure or to stabilize the post-PUM2 bound structure (**Figure 5b**). We confirmed that the mutations do disrupt RNA structures as expected by footprinting experiments (**Supp. Figure 16**). Indeed, mutations that disrupted either side of the stem in the pre-PUM2 structure resulted in a further decrease in luciferase activity (Mutations A and B, **Figure 5c**), and an increase in miR-30 binding (**Figure 5d**), indicating a shift towards post-PUM2 structure. Importantly, compensatory mutations that restored the pre-PUM2 binding structure rescued this decrease in luciferase activity and the increase in miR-30 binding (Mutation C, **Figure 5c, d**). Additionally, mutations that stabilized the post-PUM2 structure by increasing the number of GC base pairs along the stem also decreased luciferase activity and increased miR- 30 binding (Mutation D, **Figure 5c, d**). This is consistent with our conclusion that PUM2 miR-30 coregulation on LIN28A is mediated through structure.

**Figure 5.**
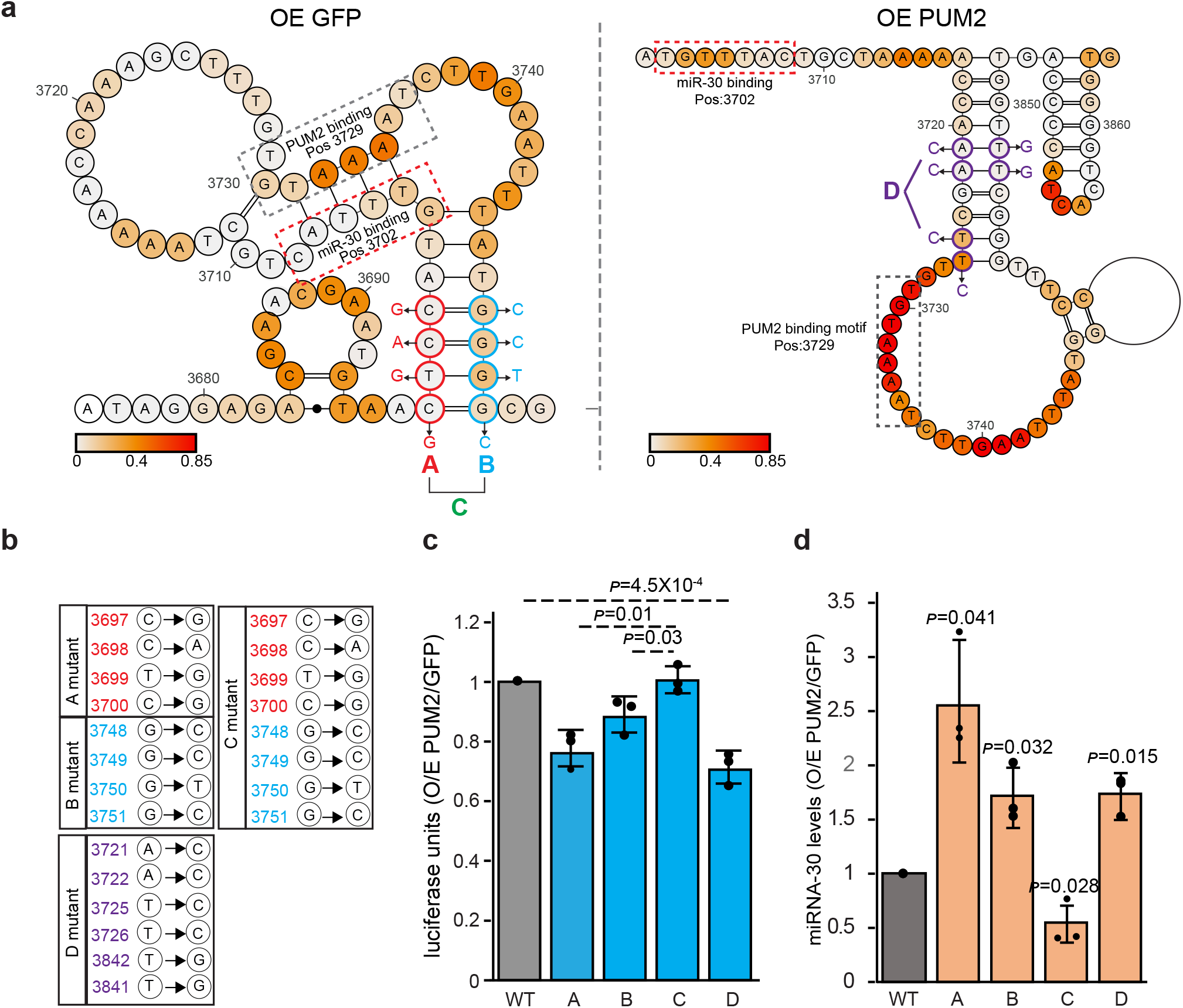
PUM2 and miR-30 co-regulates LIN28A through structure. **a**, Structural models of the pre-PUM2 binding LIN28A structure (left) and post-PUM2 binding LIN28A structure (right) based on RNA structure reactivity. PUM2 (grey dotted box) and miR-30 (red dotted box) binding sites are marked on the secondary structure. The mutational target nucleotides were labeled in bold circles, and the designed mutations were shown side by side. A, B, C and D mutants were labeled with red, cyan, green and purple color respectively. **b,** The position and sequences of designed mutants as shown in **a**. **c,** Relative luciferase units of LIN28A WT and mutants, cloned behind luciferase genes, upon overexpression of PUM2 versus GFP. *P*-values were calculated using the one-tail T-test, replicates number n=3). **d,** Ratio of miR-30 pulled down with WT LIN28A or its mutants upon PUM2 versus GFP overexpression. miR-30 levels were determined using qRT-PCR. *P*-values were calculated using the one-tail T-test, replicates number n=3.

## Discussion

Neuronal development is an extremely complex process that is extensively regulated. By using high throughput structure probing strategies, we probed the RNA structures and their dynamics during neuronal differentiation, providing an understanding of an additional layer of regulation to gene expression. One of the challenges of structure probing inside cells is that while highly reactive regions indicate the presence of single-stranded bases, low reactivity regions could either be a result of double-stranded bases or binding due other cellular factors such as RBPs and miRNA binding. Additionally, confidently identifying reactivity changing regions between two biological states can be challenging. Here, we compared different analytic strategies to identify reactivity differences by using footprinting data as the ground truth. We showed that NOISeq performs better than other methods and we believe that NOISeq can be widely used to identify reactivity changes in other cellular conditions.

Globally, we observed that RNA structures are more accessible in hESC than in differentiated cell states, reminiscent of open chromatin and pervasive transcription in hESC(Efroni et al., 2008). This suggests that RNA structures could also be “poised” for transcript regulation in hESCs depending on the direction of cellular lineage that they are heading. Recent high throughput studies show that ribosomes are a major factor for structure remodeling inside cells. Here, we show that dynamic RNA structures during differentiation are associated with miRNA, splicing, m6A modification and RBP binding, with m6A modification being enriched in CDS and associated with increased translation. This agrees with existing literature that m6A modification increases structure accessibility(Liu et al., 2015) and that modifications on CDS enhances translation.

While *in silico* modeling have suggested the presence of extensive RBP co-regulation through secondary structures(Lin and Bundschuh, 2013), only individual examples of this have been shown experimentally in a couple of cellular states, including in quiescent(Kedde et al., 2010) and hypoxic(Ray et al., 2009) conditions. Whether RBPs could coordinate gene regulation through RNA structure remodeling in a wide-spread manner in human development is still largely unknown. Our studies show that RBPs play pervasive roles in regulating RNA structures during development and modulates co-regulation networks through structure. The complexity of their regulation is demonstrated by our LIN28A-PUM2-miR-30 example, showing that PUM2 can use structure dynamics to control the miR-30 interaction with LIN28A. This study deepens our understanding of the connectivity and complexity of structure mediated gene regulation during neuronal development.

## Materials and Methods

### Cell culture

HEK293T cells were obtained from neighboring labs and cultured in DMEM with 10% FBS and 1% Pen-Strep. Human ESCs, neural stem cell induction and neuronal differentiation were performed as previously described(Wang et al., 2017). Briefly, human embryonic stem cells (H9) were cultured in mTeSR^TM^1 (#05851, stem cell technologies) medium, in the presence of 1X Supplement (#05852, stem cell technologies).

### Neuronal differentiation

Neuronal differentiation was performed as in the previous publication(Wang et al., 2017). Briefly, H9 cells were dissociated using dispase (#07923, stem cell technologies), and seeded in a new cell culture dish to 20-30% confluence. After 1-2 days, H9 cells were then treated with CHIR99021, SB431542 and Compound E in neural induction medium. We changed the culture medium every 1-2 days. After seven days, we split the cells 1:3 using accutase (# 25-058-CI, Corning) and seeded the cells on matrigel-coated plates. We then added the ROCK inhibitor (1254, Tocris) to a final concentration of 10uM when we were passaging the cells.

The derived 2 X10^5^ neural stem cells were subsequently seeded on poly-l-lysine (P4707, Sigma) and laminin (L2020, Sigma) coated 6-well plates in neural cell culture medium for 1-2 days. The cells were then cultured in neuronal differentiation medium: DMEM/F12(11330-032), 1X N2(17502-048), 1X B27(17504-044), 300ng/ml cAMP(A9501), 0.2mM vitamin C(A4544-25), 10ng/ml BDNF(450-02), 10ng/ml GDNF(450-10) for 7 days.

### RNA structure probing, icSHAPE library preparation and data analysis

icSHAPE libraries were performed as previously described(Flynn et al., 2016). Briefly, cultured cells were dissociated and transferred to a 1.5ml Eppendorf tube. The cells were then washed with PBS and treated with DMSO or 50mM NAI-N3 for 10min at 37°C with gentle rotation. RNA was then extracted using Trizol and poly(A)+ selected. We add on biotin on 0.5-1ug of poly(A)+ RNA using DIBO-biotin, before performing fragmentation, ligation, reverse transcription and PCR amplification. We performed 6-8 PCR cycles for DMSO treated samples, and 9-11 PCR cycles for NAI-N3 treated samples.

Raw sequencing reads from icSHAPE library were first trimmed to remove adaptors using cutadapt (version 1.12)(Martin, 2011), collapsed to remove PCR duplicates and trimmed to remove the leading 15-nt containing both the unique molecular identifiers and index sequences. The processed reads were then mapped to an in-house reference transcriptome using bowtie (version: 0.12.8)(Langmead and Salzberg, 2012). The in-house reference transcriptome contains the longest transcript of each gene, and was built based on the human reference genome GRCh38 and the gene annotation data from ENSEMBL version 92(2020). The SHAPE reactivity score was calculated as previously described(Spitale et al., 2015).

### Riboswitch & RiboSNich Benchmarks

1ug of each IVT RNA [6 riboswitches (with or without metabolite) and 2 riboSNiches] were refolded separately in PCR machine by heating the RNA at 90°C for 2 min, chilling on ice for 2 min, and then ramping up to 37°C at 0.1°C/s. We then incubated the RNA for another 20 min at 25°C, before treating the refolded RNA with NAI-N3 for 10 min. DMSO treatment was used as a negative control. NAI-N3 and DMSO RNA was purified using phenol:chloroform:isoamyl alcohol (25:24:1). Half of the RNA was used to run radioactive gels while the other half of the samples were used to make icSHAPE libraries. The working concentrations of the ligands for the riboswitches are: 75uM GUA (Guanine), 100uM FMN (flavin mononucleotide), 150uM SAM (S-adenosylmethionine), 100uM TPP (Thiamine pyrophosphate)

### SHAPE-MaP library preparation and data analysis

We performed SHAPE-MaP following the previously published literature (Siegfried et al., 2014) with a few modifications. Briefly, we extracted the RNAs after the cells were treated with 100mM NAI for 10min. Poly(A)+ RNA were then purified using the Poly(A)Purist™ MAG Kit (AM1922). 1ug of poly(A)+ RNA was fragmented and used to perform reverse transcription in the presence of 6mM MnCl2. This is then followed by second-strand synthesis and NEB ultra-directional RNA sequencing kit (NEB #E7760).

Raw sequencing reads from SHAPE-MaP library were first trimmed to remove adaptors using cutadapt (version 1.12)(Martin, 2011). Processed reads were then mapped to an in-house reference transcriptome using bowtie (version: 0.12.8)(Langmead and Salzberg, 2012). The mutation rate was computed as the ratio of non-reference alleles in a particular location. The reactivity was defined as

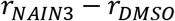

Whereby *r*_*NAIN*3_ is the mutation rate from the NAI-N3 treated library and the *r*_*DMSO*_ is the mutation rate from the DMSO treated library. Reactivities were then normalized by dividing by the mean reactivity of the top 10% of reactivities gene-wise, after reactivities above a threshold are excluded. The threshold is given as suggested by Low and Weeks(Low and Weeks, 2010).

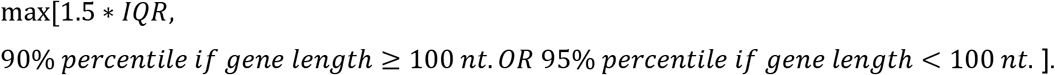

where *IQR* is the inter quantile range for the reactivities. The normalized reactivates were then used for following analysis.

### DMS-MaP-Seq library preparation and data analysis

We performed DMS-MaP-Seq following the previously published literature(Zubradt et al., 2017) with a few modifications. Briefly, total RNA was extracted after the cells were treated with 4% DMS for 5min and quenched using 30% BME. Poly(A)+ RNA were then purified using the Poly(A)Purist™ MAG Kit (AM1922). 1ug of poly(A)+ RNA was fragmented and used to perform reverse transcription by SSII in the presence of 6mM MnCl2. This is then followed by second-strand synthesis and NEB ultra-directional RNA sequencing kit (NEB #E7760).

The first 5 bases were trimmed from the raw sequencing data from DMS-MaP-Seq libraries as suggested by Zubradt et al(Zubradt et al., 2017) for SSII generated libraries. The trimmed reads were then collapsed to remove PCR duplicates using the BBMap package(Bushnell, 2014). The collapsed reads were further cleaned to remove adaptors using cutadapt (version 1.12)(Martin, 2011). The resulting clean reads were mapped to our in-house reference transcriptome using Bowtie2 (version: 2.3.5.1)(Langmead and Salzberg, 2012). The best mapping location for multiple mapping reads were selected based on the best alignment score or randomly if the alignment scores are same. The mutation rate was computed as the ratio of non-reference alleles in a particular location. Only the nucleotides whose reference allele is A or C and the mutation rate is <= 20% were considered. The raw reactivity was defined as:

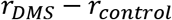

Whereby *r*_*DMS*_ is the mutation rate from the DMS treated library and the *r*_*control*_ is the mutation rate from the untreated library. To reduce noise, we filtered out the nucleotides with negative raw reactivity values. For each gene, the raw reactivity values were winsorized to 5%-95% and further scaled to [0,1]. Transcripts need to have 1) a mean depth of coverage (for A/C only) >= 100, 2) more than 50 A/C detected and 3) more than 50% of the total A/C detected to be included in analysis. The reactivity per gene is calculated as the mean reactivities across all the remaining A/C nucleotides.

### Enhanced CLIP (eCLIP) library preparation

eCLIP libraries were prepared as described in the previous publication(Van Nostrand et al., 2016b). Briefly, cells were grown on 10cm plates, washed twice with PBS and cross-linked at 254-nm UV with an energy setting of 400 mJoules/cm^2^. The cells were then harvested, lysed and sonicated. 2% of the cell lysate was used as input, and the rest of the cell lysate was incubated with anti-Flag M2 Magnetic beads (M8823-5ML) overnight at 4°C. The incubated beads mixture (IP samples) was then washed for 5 times. Input and IP RNA samples were size selected using SDS PAGE gel electrophoresis, purified and reversed transcribed into a cDNA library. The cDNA library was then amplified using PCR and deep sequenced.

### RNA pull down experiment

We pulled down LIN28A using a protocol that was previously described(Tan et al., 2014). Briefly, pcDNA-PUM2 was co-transfected with pmirglo- LIN28A-3729 or LIN28A-1485 into HEK293T cells in 6-well plates for 48 hours. The cells were then harvested and lysed. We added biotinylated probes against LIN28A into the cell lysate and allowed the probe to hybridize by incubating overnight at 4°C with continuous rotation. We then added 20ul of Dynabeads® MyOne™ Streptavidin C1(#65001) into each sample and rotated at room temperature for 45 minutes. We pulled down the RNA by placing the magnetic beads on a magnetic stand and washing the beads as previously described. Pulled down RNA is eluted and extracted from the beads using Trizol. We then performed qRT-PCR for miR-30 as previously described(Wang et al., 2012).

### In vitro RNA-protein interaction experiment

We obtained individual RNA fragments (see supplemental table) using PCR amplification and in vitro transcription. We then pooled the RNA fragments together and added 12 pmol of RNA pool into 60ul of structure probing buffer (final concentration: 50mM Tris PH7.4, 10mM Mgcl2 and 150mM Nacl). We aliquoted 20ul of RNA into 3 tubes, and refolded the RNAs in thermocycler by heating the RNA sample at 90°C for 2min, cooling down the sample at 4°C for 2min, and then slowly ramping up the sample to 37°C at 0.1°C/s. We incubated the sample for another 20min at 37°C before adding 0, 4 and 10 pmol of purified PUM2 proteins (#TP311307) into the 3 aliquots respectively and incubating for 1 hour. We then added NAI-N3 (final concentration of 50mM) or DMSO to each reaction for 10min at 37°C for structure probing. The RNAs were purified using phenol:chloroform:isoamyl alcohol (25:24:1) before being converted into a cDNA library following icSHAPE protocol(Siegfried et al., 2014).

### Luciferase assay

Target regions of PUM2 were PCR amplified and cloned into pmirGLO Dual-Luciferase Expression Vector (# 9PIE133). pcDNA-PUM2/pcDNA-GFP and target pmirGLO-candidates were then co-transfected into HEK-293T cells. After 48 hours post-transfection, cells were then lysed and luciferase assays were performed using Dual-Luciferase Reporter Assay System (#E1980).

### Identification of structure changing regions in the transcriptome

We designed a computational pipeline to identify the regions with significant structural changes. First, we split our reference transcriptome into individual windows of 20 nucleotides in length, and counted all the RT stops within the windows for each of the NAI-N3 treated libraries. We tested the optimal window size using data from two technical replicates. A window size of 20 nucleotides could obtain a per-gene correlation between technical replicates that is not significantly different from that of larger window sizes (**Supp. Figure 5e**). Next, we filtered out windows with a RT stop count of <=50 to remove poorly expressed transcripts. We then normalized the RT stop counts by calculating the RPM (reads per million mapped reads) for each window within each NAI-N3 library. Only windows with higher RPM in the NAI-N3 treated library than in the corresponding DMSO library were kept for downstream analysis. To enable data comparison between the different libraries, we performed an upper quantile normalization for each library. The RTstop counts in NAI-N3 treated library was further scaled by the ratio of RTstop counts between the corresponding DMSO libraries. To further reduce noise in our data, the windows were filtered out if (1) the average RT stop counts in the same window of the corresponding DMSO libraries was less than 20 reads; or (2) the coefficient of variation of the RT stop counts within the DMSO replicates was higher than 0.3. Finally, we applied NOISeq with biological replicates (noiseqbio function)(Tarazona et al., 2015) on the normalized RPMs of the windows in NAI-N3 treated libraries. NOIseq does not require the data to follow a particular distribution to perform differential analysis, making it suitable to analyze data from different systems and distributions. The regions with significant structural changes were defined as windows with (1) an adjusted *p*-value of not more than 0.05, and (2) having a fold-change of not less than 1.5 fold.

We validated our analysis pipeline using 6 riboSWitches and 2 riboSNiches, which are known to change structures with the presence of metabolites or mutations. To identify the structure changing regions along these RNAs, we performed gel electrophoresis in the presence and absence of metabolites or mutations. We then quantitated the relative intensity of each band on the gel using the program SAFA(DAS et al., 2005). For each riboSWitch or riboSNitch, the per-nucleotide intensity was normalized to the average intensity per aptamer as follows:

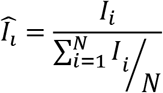

Where the 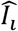 is the normalized intensity of nucleotide *i*; *I_i_* is the raw intensity of nucleotide *i* from SAFA; *N* is length of the aptamer used to run gel. Based on the assumption that the majority of the nucleotides in a riboswitch and riboSnitch will not change its secondary structure in the presence of metabolites or mutations, the distribution of *N*_*i*_ before and after structural change should be approximately the same. The difference of *N*_*i*_ will hence follow a normal distribution. Therefore, we calculated a Z-score, *Z*_*i*_, for each nucleotide as follows:

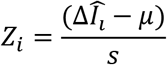

Where the 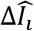 is the difference of 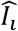 before and after structural changes; *μ* is the average of 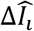; and *s* is the standard deviation of 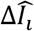. A nucleotide is defined as having a significant secondary structure change when the corresponding *Z*_*i*_ is >=1.96 or <=-1.96.

We next performed our pipeline to identify the structural changed regions in riboswitches and riboSNitches using icSHAPE data. All the cutoff and filters were applied as described above, except that the RTstop count per nucleotide is used as input, and the minimum RTstop count was set to 2.5 (based on the RTstop cutoff of 50 in a 20-nt window). This is due to the short length of riboswitches and riboSNitches used in footprinting and SAFA analysis. To benchmark the performance of our pipeline, we also tested other metrics to quantify the structural changes: (1) “delta” by taking the absolute difference between SHAPE reactivities per nucleotide. (2) “delta- win” by taking the absolute difference between average SHAPE reactivities in a 10-nt sliding windows with a step size of 1-nt. (3) “T-test” by considering the p-value from a t-test using SHAPE reactivities in a 10-nt sliding windows with a step size of 1-nt. The method is adopted from Shi et al.(Shi et al., 2020a) (Shi et al., 2020b). (4) “correlation” by taking the pearson correlation between SHAPE reactivities in a 10-nt sliding windows with a step size of 1-nt. (5) “diff-BUMHMM” by taking the posterior probability of modification changes. The method is adopted from Marangio et al.(Marangio et al., 2020). An AUC score was obtained for each method by comparing the variables described above with the structure changed nucleotides in riboswitches and riboSNitches from SAFA analysis. For our pipeline (termed as NOISeqRT), the p-values from NOISeq were used to obtain AUC score. Among all of the methods tested, NOISeq obtained the highest AUC score of 0.777 (**Supp. Figure 5c**). In addition, adding an absolute fold changes filter of 1.5 further increases the AUC score for NOISeq to 0.824.

### Filtering of windows associated with alternative splicing

As we are using a redundant reference transcriptome whereby only the longest transcript for each gene is included, differential splicing of the same gene might result in windows being artificially called to have significant structural changes. In order to filter these windows out, we identified the genes with different transcripts between the two time points. We first used DRIMseq to find the genes with significant differential transcript usage. Then, we identified the genes with dominant isoform changes based on TPM calculations using Salmon. The genes with both differential transcript usage by DRIMseq and dominant isoform changes by RPM were defined as genes with “significant dominant isoform changes”. The windows within the genes were removed for the downstream analysis. For the genes that show significant differences in isoform abundance between different time points using DRIMseq, but had the same isoform as the dominant isoform across time points, we filtered out the windows located +/- 40bp to the splice sites of the genes.

### Ribosome profiling library preparation and analysis

Ribosome profiling was performed by following Illumina TruSeq Ribo Profile (mammalian) Library Prep kit (RPHMR12126). The raw sequencing reads from Ribosome profiling library were first processed using cutadapt (version 1.12)(Martin, 2011) to remove adaptor sequences and reads with low quality or short length. The processed reads were then mapped to the reference genome (UCSC hg19 with annotations from GENCODE v.19) using program STAR (version 2.5.0c)(Dobin et al., 2013). The expression level per gene was quantified by Cuffdiff (version 2.2.1)(Trapnell et al., 2013). The gene translation efficiency (TE) was defined as the log2 value of the ratio between RPKM of the footprinting library and RNA sequencing library. The genes with significant differences between TE were identified using NOISeq optimized for the use on biological replicates(Tarazona et al., 2015).

### RNA sequencing library preparation and analysis

RNA sequencing library was performed using the NEBNext® Ultra II Directional RNA Library Prep Kit for Illumina (NEB #E7760L), following manufacturer’s instructions. The raw sequencing data from RNAseq library were first processed using cutadapt (version 1.12)(Martin, 2011) to remove adaptor sequences and reads of low quality or short length. The processed reads were then mapped to the reference genome (UCSC hg19 with annotations from GENCODE v.19) using STAR (version 2.5.0c)(Dobin et al., 2013). The differentially expressed genes were identified using Cuffdiff (version 2.2.1)(Trapnell et al., 2013). In addition, we also quantified the level of transcripts using the program Salmon (version 0.11.2)(Patro et al., 2017). Transcript annotations were obtained from GRCh38 from ENSEMBL (version 92)(2020). The differentially expressed transcripts were identified using DRIMseq from Bioconductor(Nowicka and Robinson, 2016).

### RNA decay library preparation and analysis

To determine the half-life of cellular RNAs, we stopped transcription by treating cells with 5µM of Actinomycin D (A1410, Sigma) for 0, 1, 2, 4 and 8 hours by following previous report(Mizrahi et al., 2018). We then extracted the RNA using Trizol, used 1-2ng RNA for reverse transcription and performed 8 cycles of PCR amplification. The derived PCR products were then used to make into a sequencing library using the Nextera XT DNA Library Prep kit (FC-131-1096).

The raw sequencing reads from mRNA decay library were mapped to our reference genome (GRCh38) using STAR (version 2.5.0c)(Dobin et al., 2013). The gene annotation data was collected from ENSEMBL(2020). The RPM values of each gene were quantified using featureCounts (version 1.6.3)(Liao et al., 2014), and then used to estimate mRNA half-life. We used the mean RPM values of a few house-keeping genes (GAPDH, ACTB, PGK1, PPIA, RPL13A, RPLP0) that are known to have consistent expression upon Actinomycin D treatment as reference, and normalized the expression levels of other genes to them across different time points post treatment (0h, 1h, 2h, 4h and 8h after treatment). The normalized expression levels were used to fit into an exponential decay model for each gene to calculate half-life:

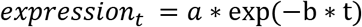

Where ***a*** is the expected expression before treatment, ***b*** is the decay constant and ***t*** is time in hours after treatment. A minimum correlation of 0.75 between fitted and observed gene levels was used to filter out genes with poor fit. Genes with significant differences in estimated half-lives between two time points were identified using NOISeq(Tarazona et al., 2015).

### RBP enrichment and PrismNet analysis

The binding sites of 183 different RNA binding proteins (RBP) determined by eCLIP(Van Nostrand et al., 2016a) were downloaded from ENCODE. We then overlapped the binding sites of each RBP with reactivity changing and non-changing regions. The enrichment of RBP in the reactivity changing versus non-changing regions were tested using hypergeometric tests. The resulting *p*- values were FDR corrected using the bonferroni method. RBPs with an adjusted *p*-value of smaller than or equal to 0.01 are considered as enriched.

Independent of eCLIP, we also developed a de novo approach using sequence motifs collected from the ATtRACT database(Giudice et al., 2016) to identify RBP binding motifs that are enriched in the reactivity changing regions. First, we identified 200 different 6-mer sequences enriched in the reactivity changing regions using hypergeometric test with bonferroni correction (adjusted *p*- value cutoff = 0.01). Then the enriched 6-mers were compared to the known regulatory motif sequences using pairwise local sequence alignment. A motif was defined as enriched when its sequence is similar to the 6-mer with at most 1 base difference. The enriched RBP by motif were defined as the RBPs with at least 1 motif sequence enriched in the structure changed regions. Last, we identified 10 RBPs that are significantly enriched in structure changing regions by using both the binding sites and motif sequence enrichment analysis. In addition to eCLIP and Motif based methods, we also used the PrismNet predicted RBP binding sites to perform the RBP enrichment analysis in the structural changing regions. The predicted RBP binding sites in H9 cells were adopted from Sun et.al(Sun et al., 2021)

### miRNA enrichment analysis

The potential miRNA binding sites are obtained from the predicted non-conserved miRNA sites in TargetScan release 7.2(Agarwal et al., 2015). The miRNA used for analysis were the highly expressed miRNA in hESC and NPC (top 100 RPM using small RNA sequencing from both developmental stages). The enrichment of miRNA near the RBP binding sites with reactivity changing regions were computed using hypergeometric test.

### Analysis of m6A binding sites

m6A binding sites in hESC were obtained from previous literature by Batista et al.(Batista et al., 2014). The binding sites are presented as 100-nucleotide regions in genomic coordinates. We converted the genomic coordinates into transcriptomic coordinates by matching the sequences.

### RNA structure modelling

Secondary structures of LIN28A were predicted by incorporating *in vivo* icSHAPE data into the program RNAstructure(Deigan et al., 2009). Use of raw icSHAPE data in structure modeling was precluded, as the discontinuous nature of the data caused divergence in partition function calculations. Hence, we proceeded to smooth the raw icSHAPE data in a 5 nucleotide window moving average before incorporating it into secondary structure modeling. We used default parameters for shape intercept (-0.6 kcal mol^-1^) and shape slope (1.8 kcal mol^-1^) respectively. Separate partition functions were calculated for LIN28A regions 1390-2000, 2200-2600 and 3600- 4014 after evaluating the likely structure boundaries of a full-length LIN28A model. After obtaining partition functions, pairing probabilities for all bases were extracted and Shannon entropy calculated for each position. Concurrently, maximum likelihood structures were calculated based on the partition functions. VARNA(Darty et al., 2009) was used for visualization and moving average icSHAPE scores were added as a color map.

Separate structural models were constructed from icSHAPE data obtained under conditions of overexpression GFP control and PUM2 respectively and changes in LIN28A structures under these conditions are shown in **Figure 5a**. Mutants for testing PUM2 binding were subsequently designed based on these structure models (**Figure 5a, b**).

## Acknowledgements

We thank members of the Wan lab for helpful discussions. YW is supported by funding from A*STAR Investigatorship, EMBO Young Investigatorship and CIFAR Azrieli global scholar fellowship.

## Author contributions

Y. Wan conceived the project. Y. Wan, JX. Wang designed the experiments. JX.Wang and W.Tan performed all the experiments. T. Zhang, M. Wen and Y. Shen performed the computational analysis. R. Huber did the structure modeling of LIN28A. Y. Wan and JX. Wang organized and wrote the paper with all other authors.

## Supplementary Figure Legends

**Supplementary Figure 1.**
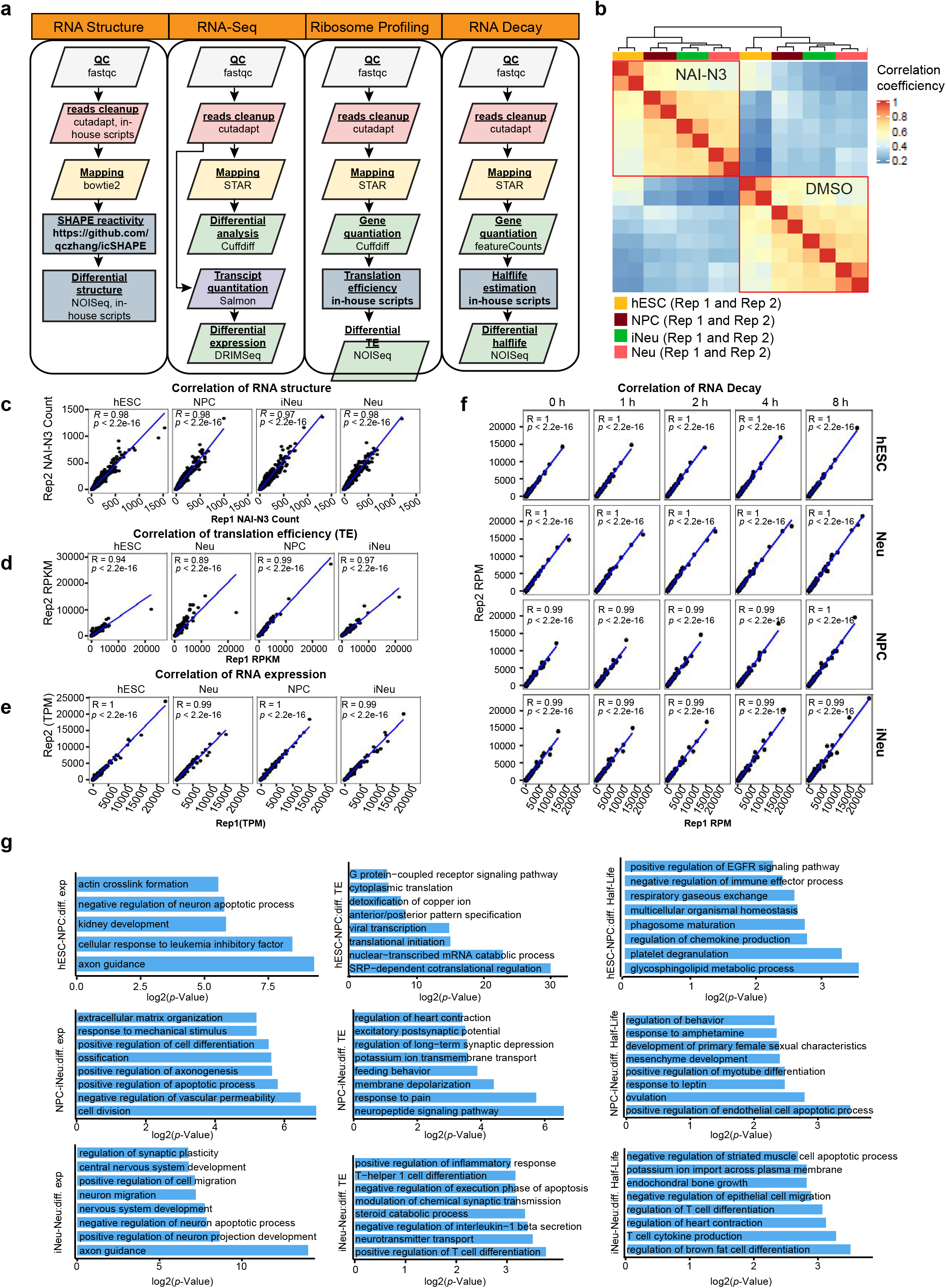
Data quality of high throughput libraries. **a,** Workflow of the multi-omics analysis framework. **b,** Heatmap of correlation between the biological replicates of hESC, NPC, iNeu and Neu treated with both DMSO and NAI-N3. **c**, Correlation of RT stop counts between 2 biological replicates of icSHAPE libraries for hESC, NPC, iNeu and Neu, from left to right. **d,** Correlation of RPKM values per gene between 2 biological replicates of ribosomal profiling libraries for hESC, NPC, iNeu and Neu, from left to right. **e,** Correlation of TPM values per gene between 2 biological replicates of RNA sequencing libraries for hESC, NPC, iNeu and Neu, from left to right. **f,** Correlation of RPM values per gene between 2 biological replicates of RNA decay libraries at different time points post treatment (left to right), and from hESC, NPC, iNeu and Neu (top to bottom). The Pearson correlation (*R*) and the *p*-value are shown inside the subplots **c-f**. **g,** GO enrichment analysis of genes with differential expression, translation efficiency and RNA half-life between neighbor time points.

**Supplementary Figure 2.**
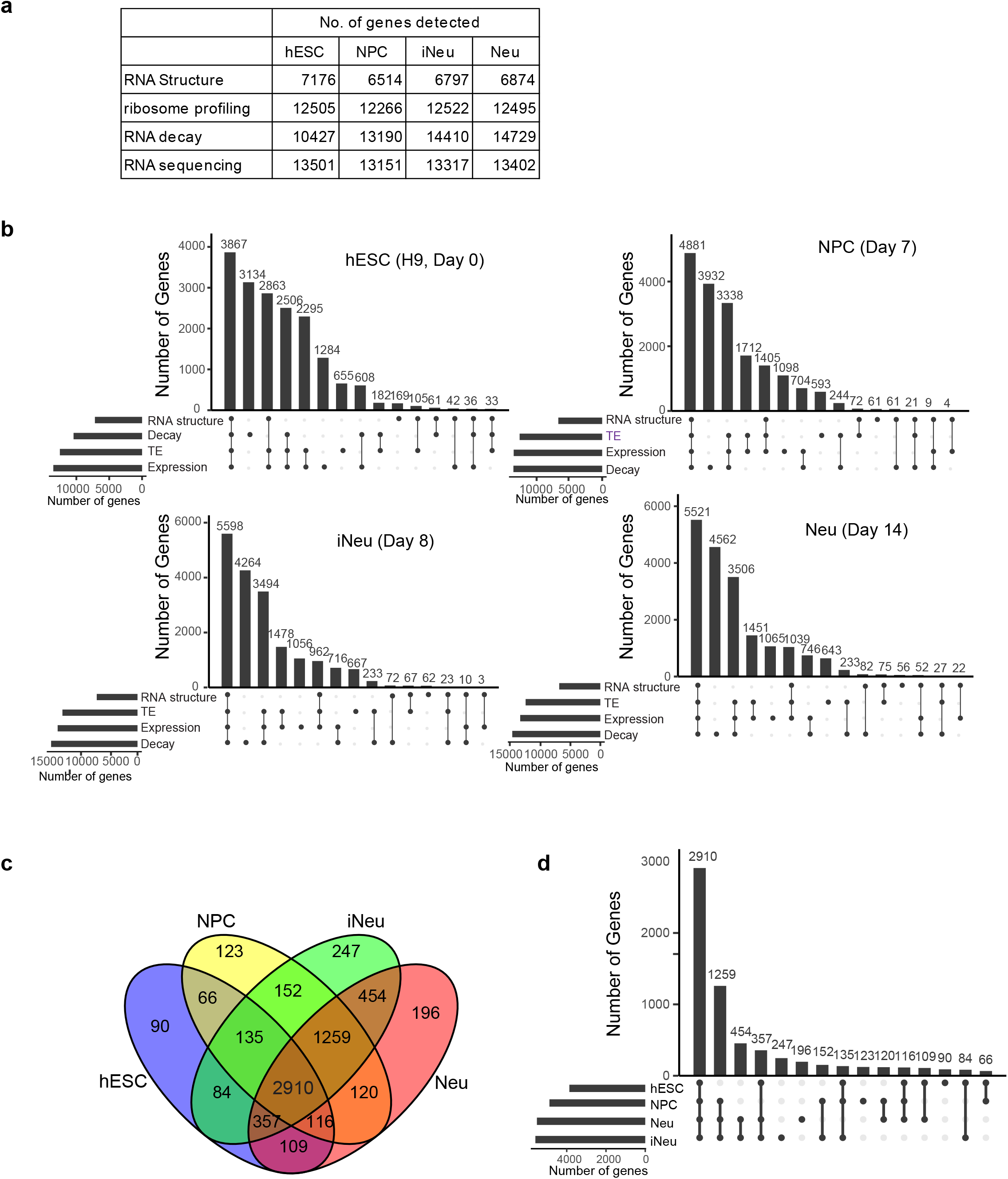
Summary of detected genes by different methods. **a,** Number of genes detected at each stage of neuronal differentiation by icSHAPE (RNA structure), ribosomal profiling, RNA decay and RNA sequencing, respectively. **b,** Upset plot showing the number of shared genes detected by different methods for hESC, NPC, iNeu and Neu, respectively. **c,** Venn diagram showing the overlapping of genes detected by all 4 methods in all stages of differentiation. **d,** Upset plot showing the number of genes detected by all 4 methods in all stages during differentiation.

**Supplementary Figure 3.**
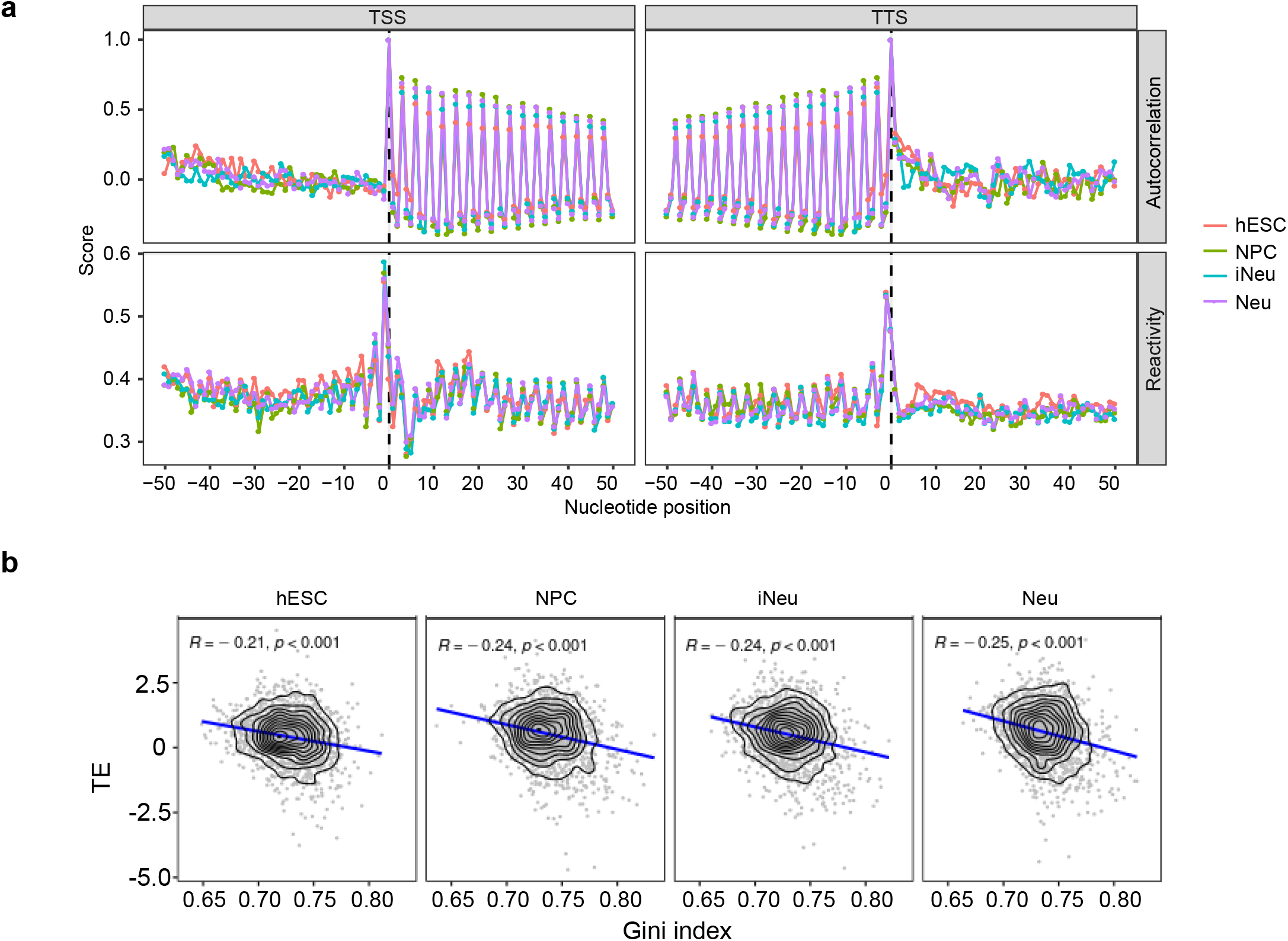
Global features of RNA structures using icSHAPE. **a,** *Bottom*: Metagene analysis of icSHAPE reactivity centered on translation start and stop sites showed a three-nucleotide periodicity in the coding region across all differentiation stages. *Top*: Autocorrelation of the reactivity in the 4 stages from hESC to Neu (Red-hESC, green-NPC, light blue-iNeu and purple-Neu). **b**, Correlation between the Gini index of reactivity from icSHAPE and TE calculated from ribosome profiling for hESC, NPC, iNeu and Neu, from left to right. The Pearson correlation (*R*) and the *p*-value are shown inside the plots.

**Supplementary Figure 4.**
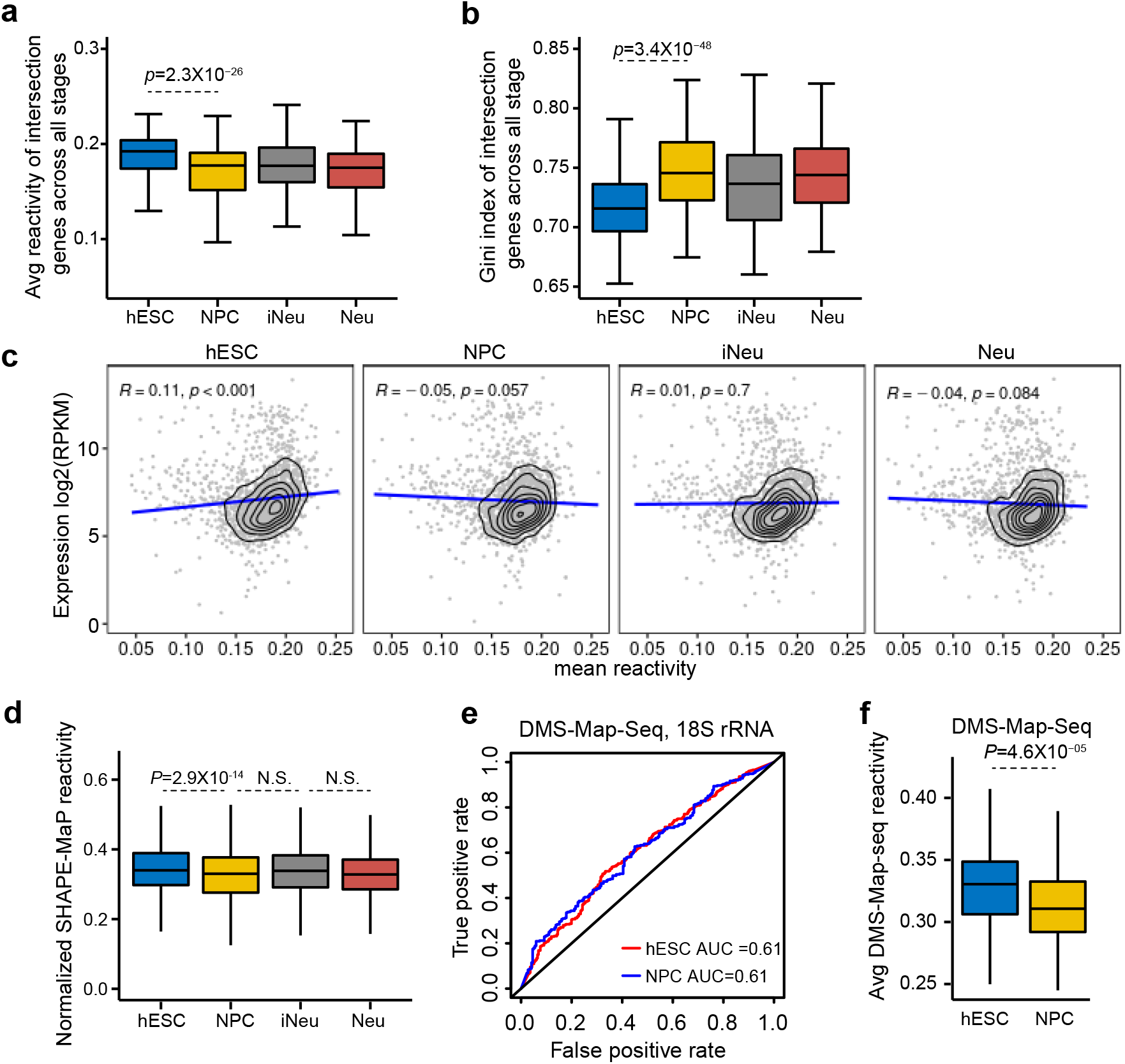
Genes in hESC are more accessible than in NPC. a, b, Boxplots showing the distribution of the mean icSHAPE reactivities (**a**) and Gini index (**b**) per gene for hESC, NPC, iNeu and Neu. The genes used here are detected by icSHAPE in all 4 stages. *P*- value is calculated using Student’s T- Test. **c,** Correlation between gene expression levels and the mean icSHAPE reactivity in hESC, NPC, iNeu and Neu from left to right. The Pearson correlation (*R*) and the *p*-value are shown inside the plots. **d,** Boxplot of the average reactivities per gene obtained by SHAPE-MaP in all 4 stages. **e,** The AUC-ROC curves of the normalized reactivities obtained by DMS-MaP-Seq in hESC and NPC cells using the 18S rRNA. The AUC-ROC values are labeled in legend. **f,** Boxplot of the average reactivities per gene obtained by DMS-MaP-Seq in hESC and NPC cells.

**Supplementary Figure 5.**
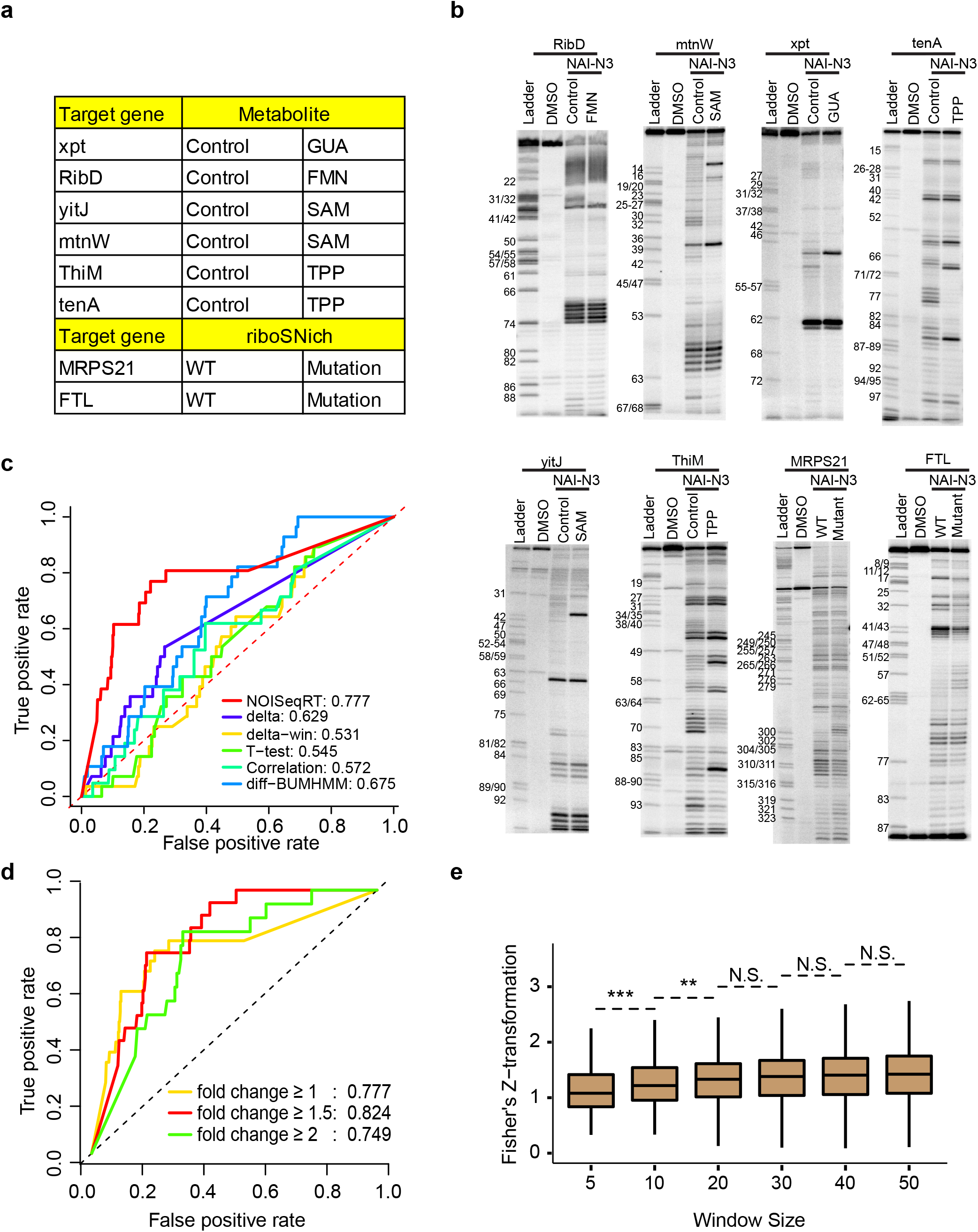
Benchmarking of icSHAPE analytic methods using riboswitches and riboSNitches. **a,** Table of riboswitches and riboSNitches used for benchmarking. **b,** Footprinting images of riboswitches and riboSNitches. For each gel, we are displaying G ladder (lane 1), DMSO treated RNA (lane 2), NAI-N3 treated RNA without ligand (riboswitch) or WT (riboSNitch) (lane 3) and NAI-N3 treated RNA with ligand (riboswitch) or Mutant (riboSNitch) (lane 4). **c,** The AUC-ROC curves of the different methods (NOISeqRT in red, delta in blue, delta- window in black, T-test in green, correlation in cyan, and diff-BUMHMM in light blue) used for calculating reactivity changes. The AUC-ROC scores are labeled in legend box. **d,** The AUC-ROC curves of different fold-change cutoffs (1X is in blue, 1.5X is in red, and 2X is in green) used in the NOISeqRT method. The AUC-ROC scores are labeled in legend box. **e,** Boxplots of per gene correlations between hESC replicates using different window sizes ranging from 5 to 50 nucleotides from left to right.

**Supplementary Figure 6.**
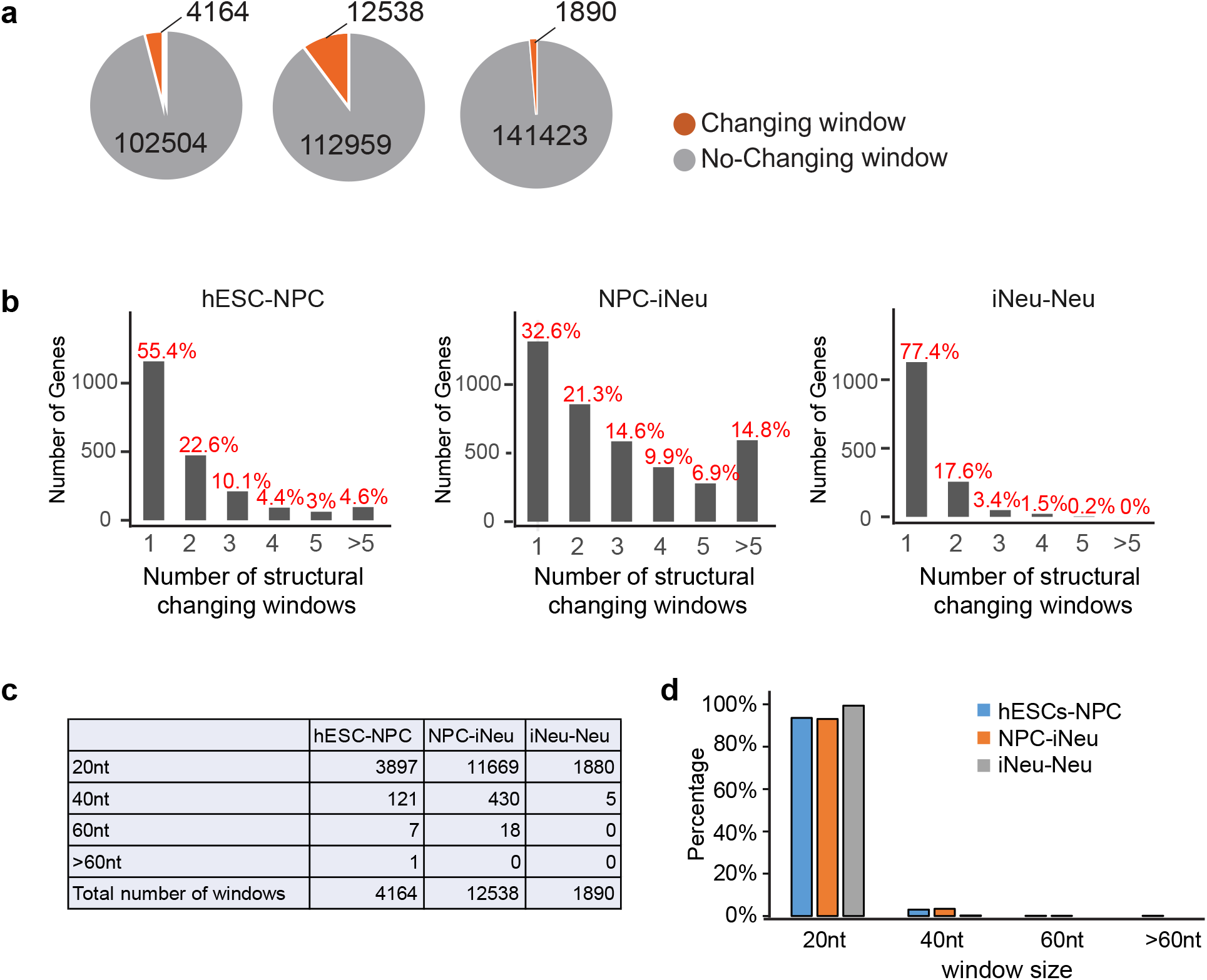
Summary of the significantly structural changing windows. **a**, Pie charts showing the numbers of windows with (orange) and without (grey) significant reactivity changes during neuronal differentiation. **b,** Histogram showing the number of genes with 1 or more significant structural changing windows during hESC-NPC (left), NPC-iNeu (middle) and iNeu-Neu (right). **c,** Table showing the number of significantly structural changing regions with different sizes after merging neighboring significant changing windows in hESC-NPC, NPC-iNeu and iNeu-Neu. **d,** Bar plot showing the percentage of significantly structural changing regions with the size of 20nt, 40nt, 60nt and more during hESCs-NPC (blue), NPC-iNeu (orange) and iNeu- Neu (grey).

**Supplementary Figure 7.**
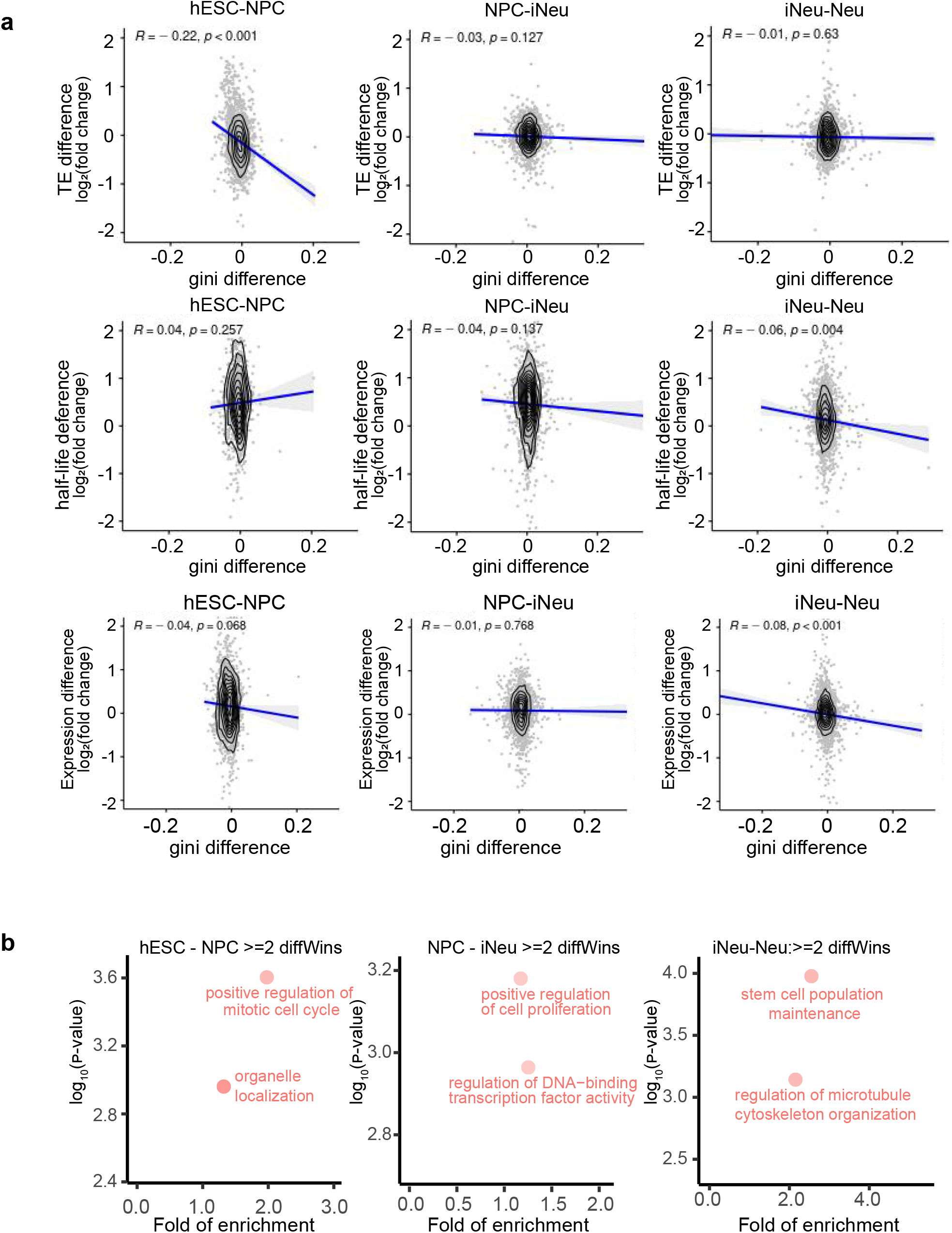
The association of structural changes with other methods. **a,** Correlation of changes in Gini index with changes in TE (top row), half-life (middle row), and gene expression (bottom row) during hESCs-NPC, NPC-iNeu and iNeu-Neu differentiation, from left to right. **b,** Enriched biological processes obtained by GO analysis for the genes with >=2 significant structural changing regions during hESCs-NPC, NPC-iNeu and iNeu-Neu differentiation from left to right.

**Supplementary Figure 8.**
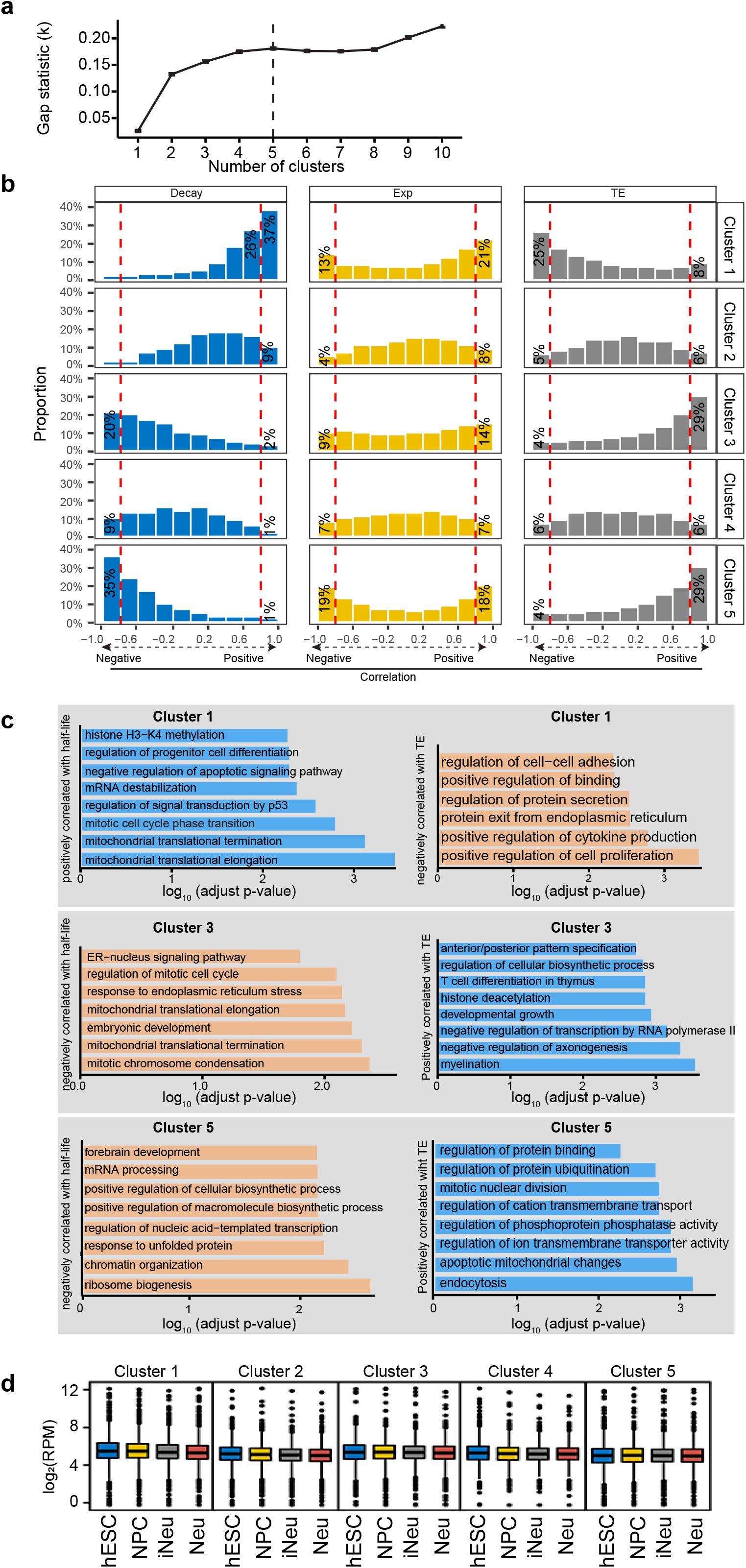
Clustering of significant structural changing windows. **a,** Line chart showing the number of optimal number of K means clusters predicted using Gap statistics. The Y-axis indicates the Gap statistic and the X-axis indicate the number of clusters. **b,** Histogram showing correlations between normalized RT stop counts by icSHAPE and half-life estimated from RNA decay (blue), gene expression level from RNA sequencing (yellow) or translation efficiency from ribosomal profiling (grey) for cluster 1 to 5 (top to bottom panels). The red dashed lines indicate the correlation values at -0.8 and 0.8 in each subplot. **c,** Barplot showing *p*-values of enriched biological processes from GO analysis using the corresponding genes of windows from cluster 1 (top row), 3 (middle row) and 5 (bottom row) with either positive correlation (colored in blue, R>=0.8) or negative correlation (colored in orange, R<=-0.8) with half-life (Decay) or TE (Translation Efficiency). **d,** Boxplot showing the expression levels of genes from clusters 1 to 5 from left to right for hESCs (blue), NPC (yellow), iNeu (grey) and Neu (red).

**Supplementary Figure 9.**
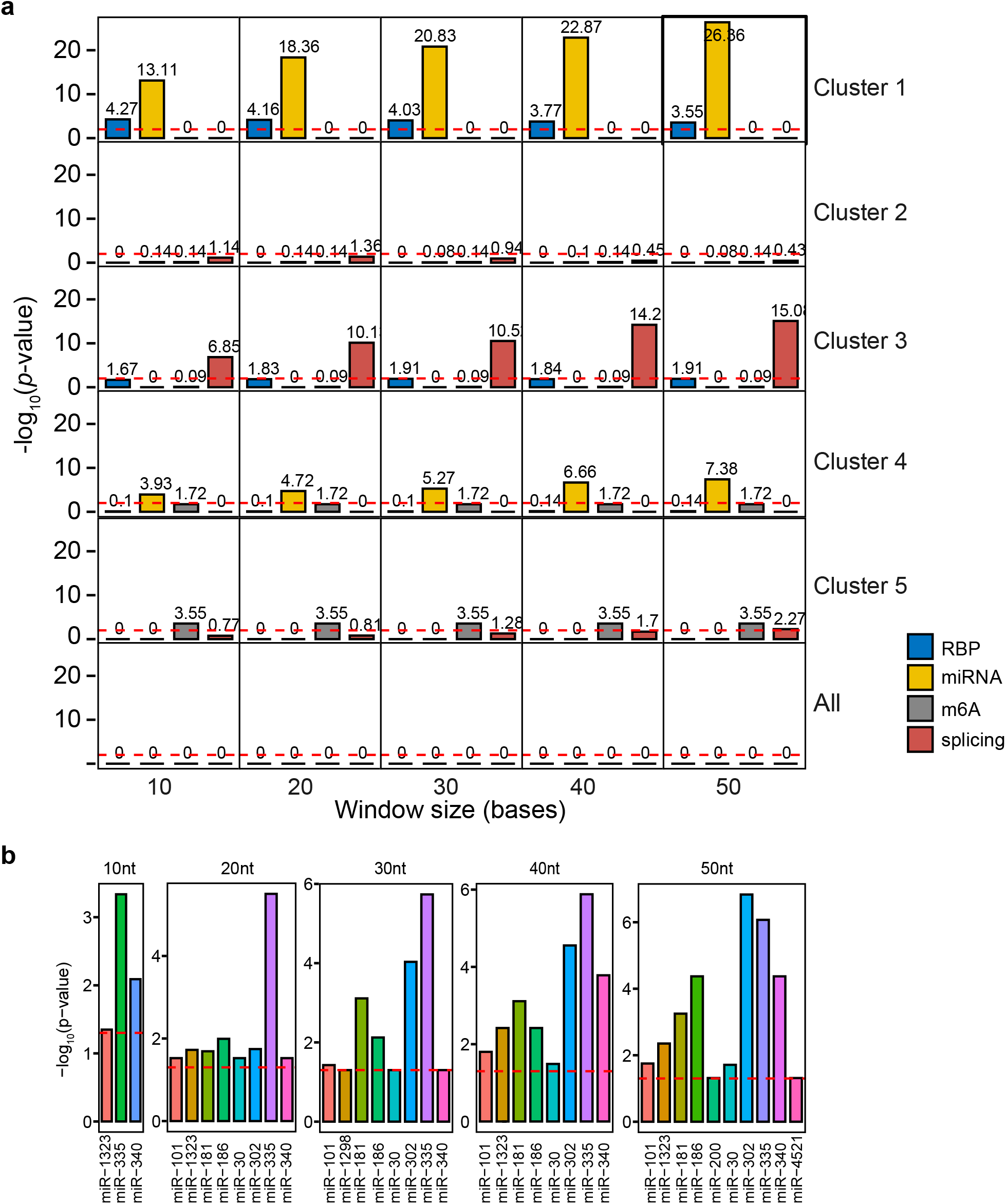
Overlap of reactivity changing windows with cellular regulators. **a**, *P*-value of enrichments of cellular factor binding sites in reactivity changing windows in clusters 1-5 and all clusters (from top to bottom). We calculated the enrichment of predicted binding sites of RBPs (blue), miRNAs (yellow), m6A (grey) and splicing sites (red) that overlap with 10-50 nucleotides upstream and downstream from the reactivity changing region. *P*-value is calculated using the binomial test. The expected proportion of overlap for each regulator is calculated from all overlapping proportions of different regulators to all windows in 5 clusters (bottom row). **b**, Barcharts showing significantly enriched miRNAs in cluster 1 using with varying flanking regions from 10 to 50 nucleotides (from left to right) of reactivity changing regions.

**Supplementary Figure 10.**
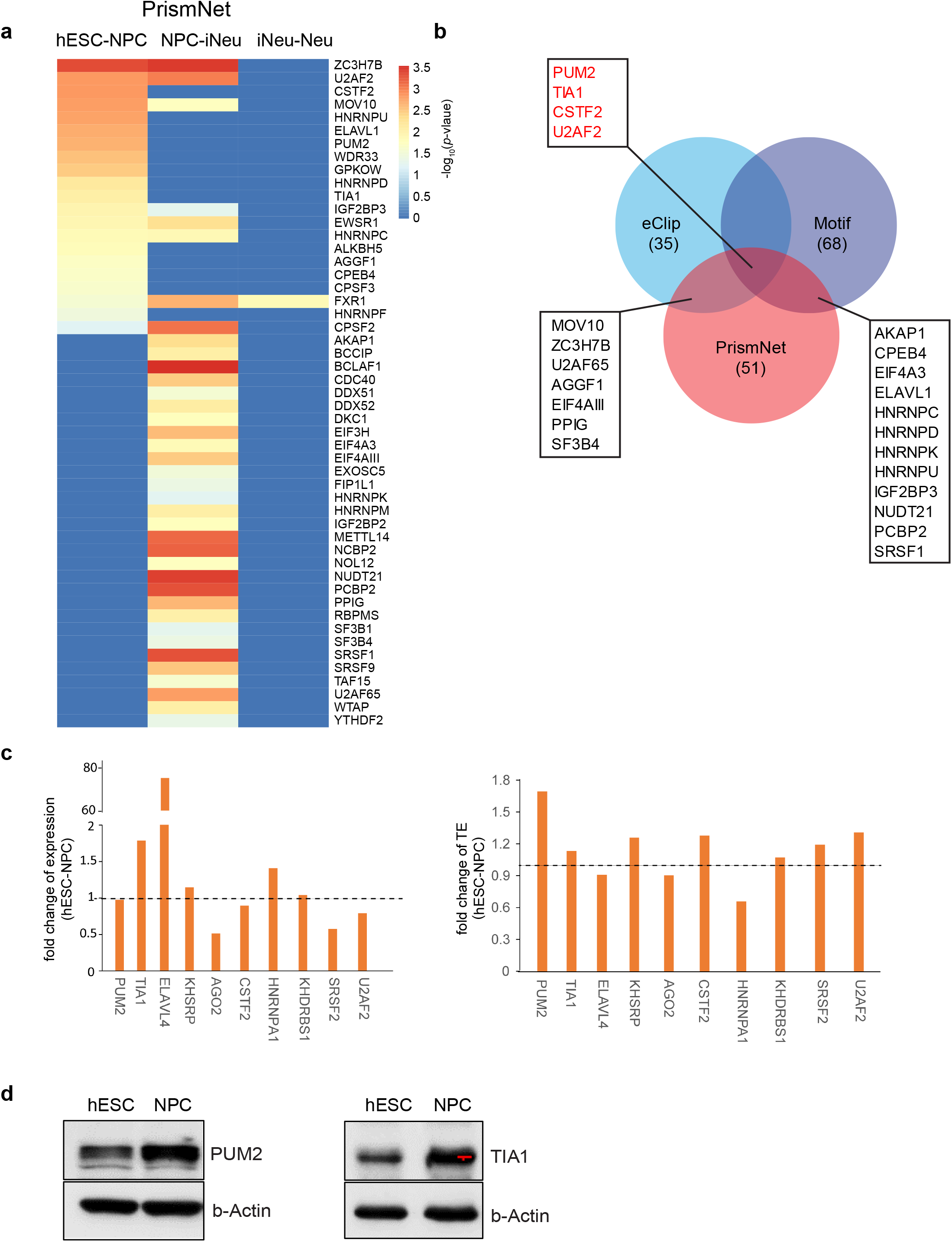
Enrichment of RBPs in significant structure changing windows. **a,** Heatmap of *p*-values (transformed by -log10) of RBP binding sites enrichments in significant structural changing windows during hESC-NPC, NPC-iNeu and iNeu-Neu. **b,** Venn diagram of the significantly enriched RBPs in structural changing windows using binding sites predicted by eCLIP, by RBP binding motifs and by PrismNet. **c,** Barcharts showing the fold changes of expression (left) and translation efficiency (right) of the 10 enriched RBPs from hESCs to NPC(Figure 3b). **d,** Western blot experiments showing the protein levels of PUM2 (left) and TIA1 (right) in hESC and NPCs. Beta-actin is used as loading control.

**Supplementary Figure 11.**
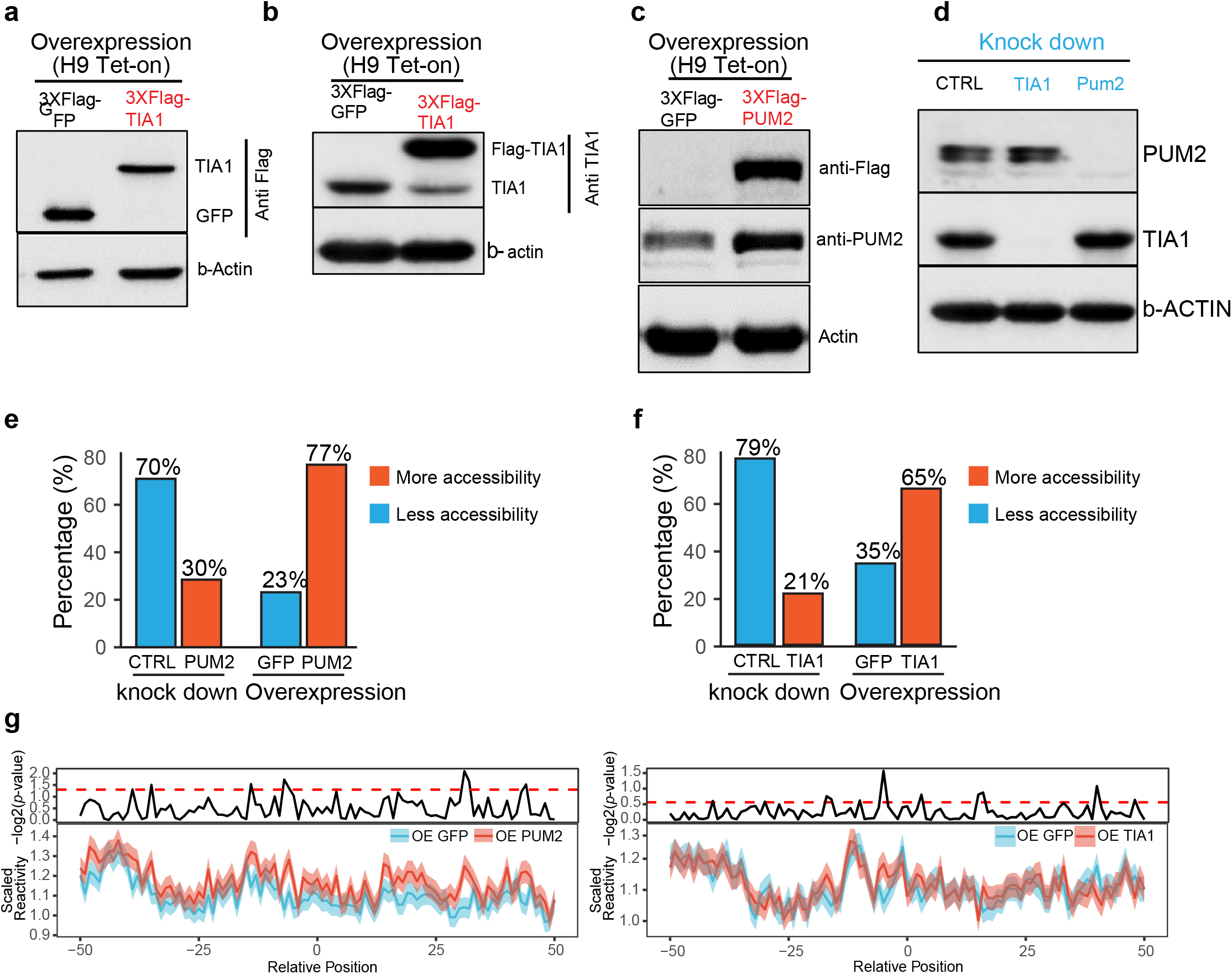
Overexpression of PUM2 and TIA1 in hESC. **a, b**, Western blot experiments using antibodies against FLAG tag (**a**) or TIA1 (**b**) when cells are over-expressed with 3X FLAG-GFP or 3X FLAG TIA1. B-actin is used as loading control. **c**, Western blot experiments using antibodies against FLAG tag (top) or PUM2 (middle) when cells are over-expressed with 3X FLAG-GFP or 3X FLAG PUM2. B-actin is used as loading control. **d,** Western blot experiments using antibodies against PUM2 and TIA1 in control, PUM2 knockdown cells and TIA knockdown cells. B-actin is used as loading control. **e,** Barcharts showing the percentages of PUM2-targeting windows becoming significantly more accessible (orange) or less accessible (blue) in knockdown (KD) and overexpression (OE) of PUM2. **f,** Barcharts showing the percentages of TIA1-targeting windows becoming significantly more accessible (orange) or less accessible (blue) in knockdown (KD) and overexpression (OE) of TIA1. **g,** Metagene analysis of normalized icSHAPE reactivity around predicted binding sites of PUM2 (left) or TIA1 (right) by PrismNet before and after PUM2 (left) or TIA1 (right) overexpression. *P*-values on top of plot were calculated per nucleotide using the single-sided T-test. The center of each PrismNet predicted binding sites were treated as relative position =0.

**Supplementary Figure 12.**
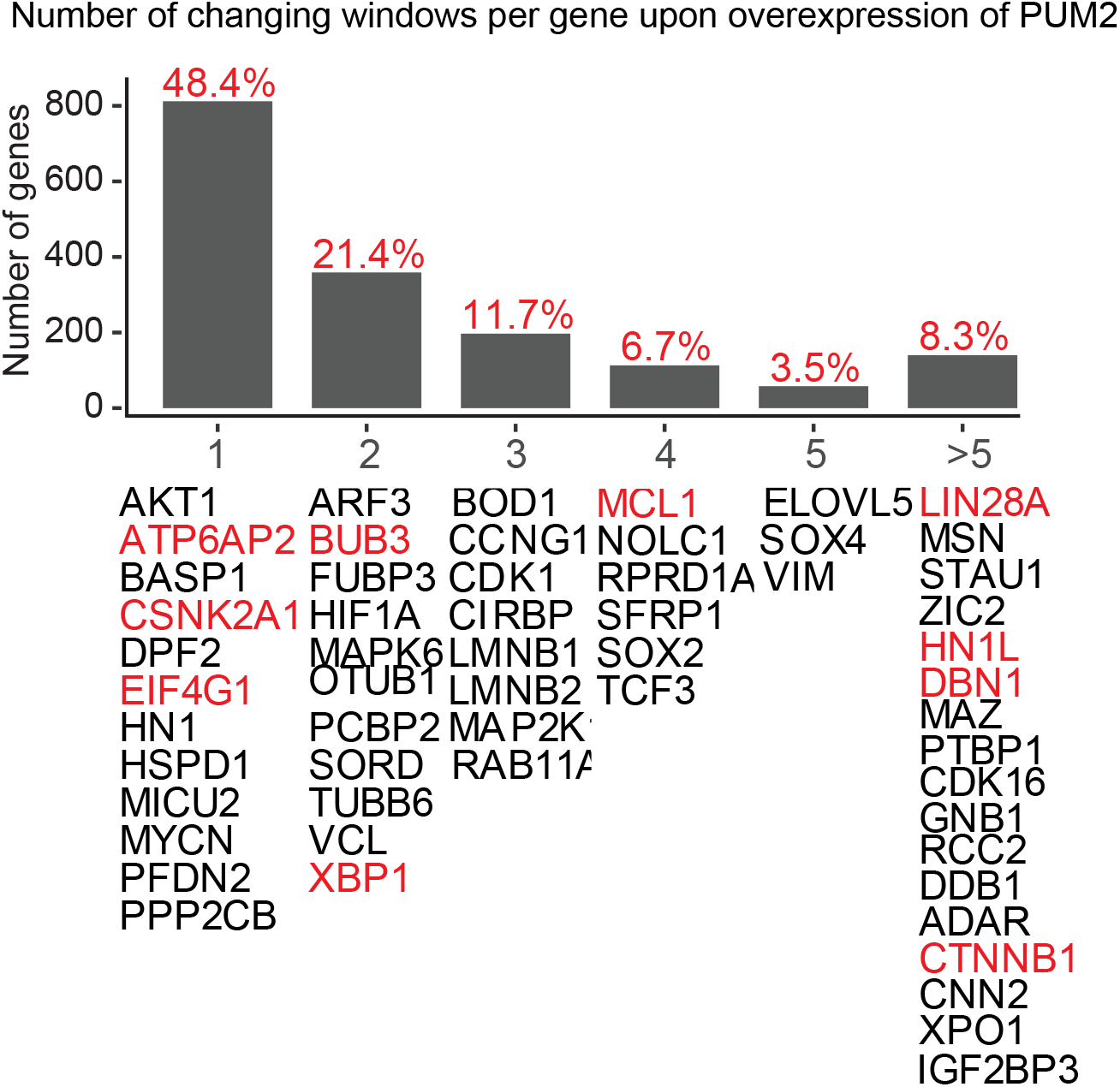
Number of changing windows per gene upon PUM2 over- expression. The distribution of number of structural changing windows per gene upon overexpression of PUM2 in hESC. Representative genes are shown below the barchart. The genes selected for luciferase assays are highlighted in red.

**Supplementary Figure 13.**
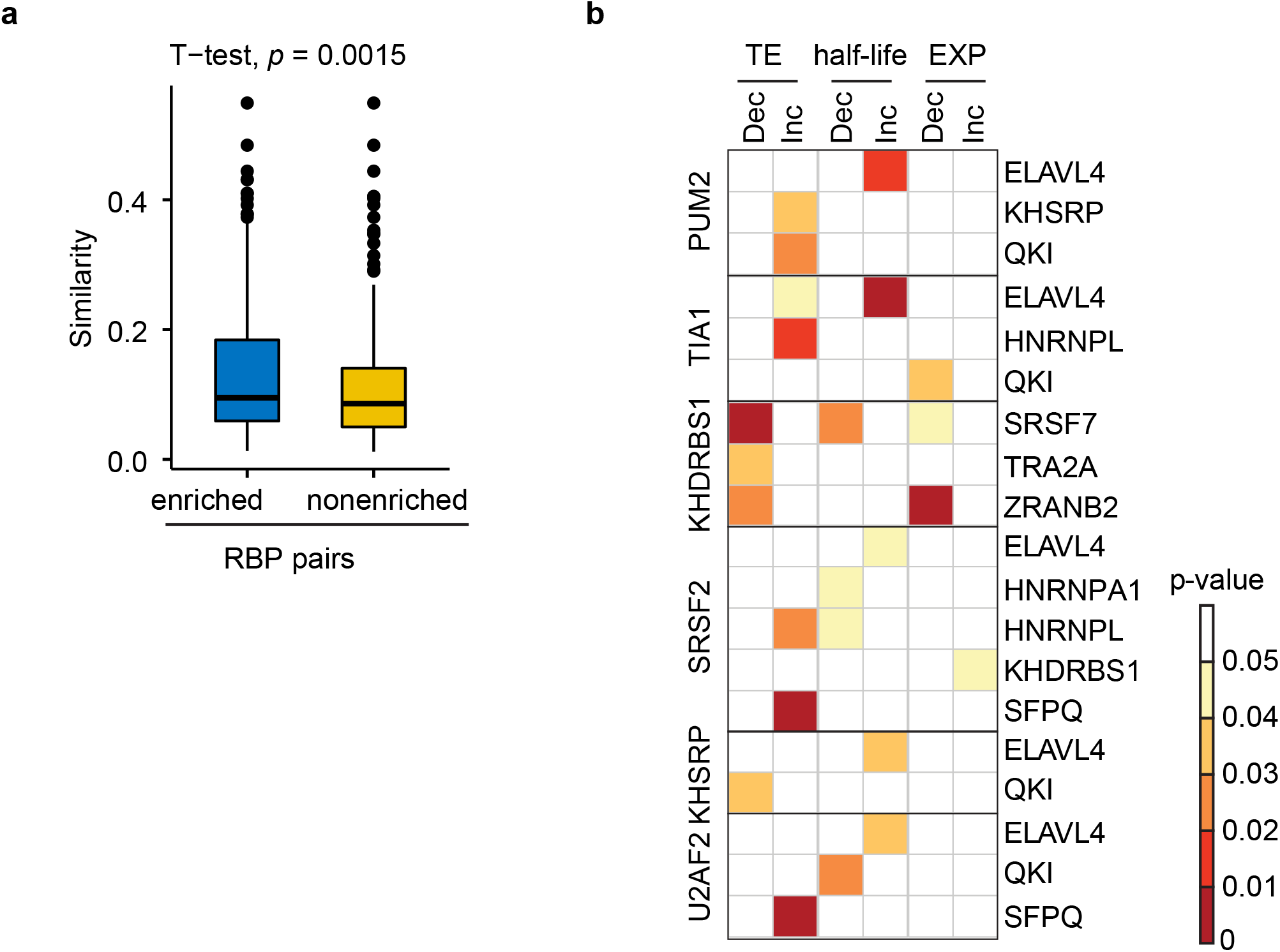
Regulation of RBP-RBP and RBP-miRNA interactions through structure. **a,** Boxplot showing the Jaccard similarity of the RBP targeting windows between enriched RBP pairs (blue) and non-enriched RBP pairs (yellow). The enriched and non-enriched RBP pairs are shown in Figure 4A. **b,** Heatmap showing changes in gene regulation (translation, decay and gene expression) of targets of RBP pairs when there are structure changes versus when there are no structure changes. *P*-value of enrichment is calculated using T-test. ‘Dec’ represents decrease in translation efficiency, half-life or expression, ‘Inc’ represents increase in TE, half-life or expression. *P*-value of enrichment is calculated using hypergeometric test.

**Supplementary Figure 14.**
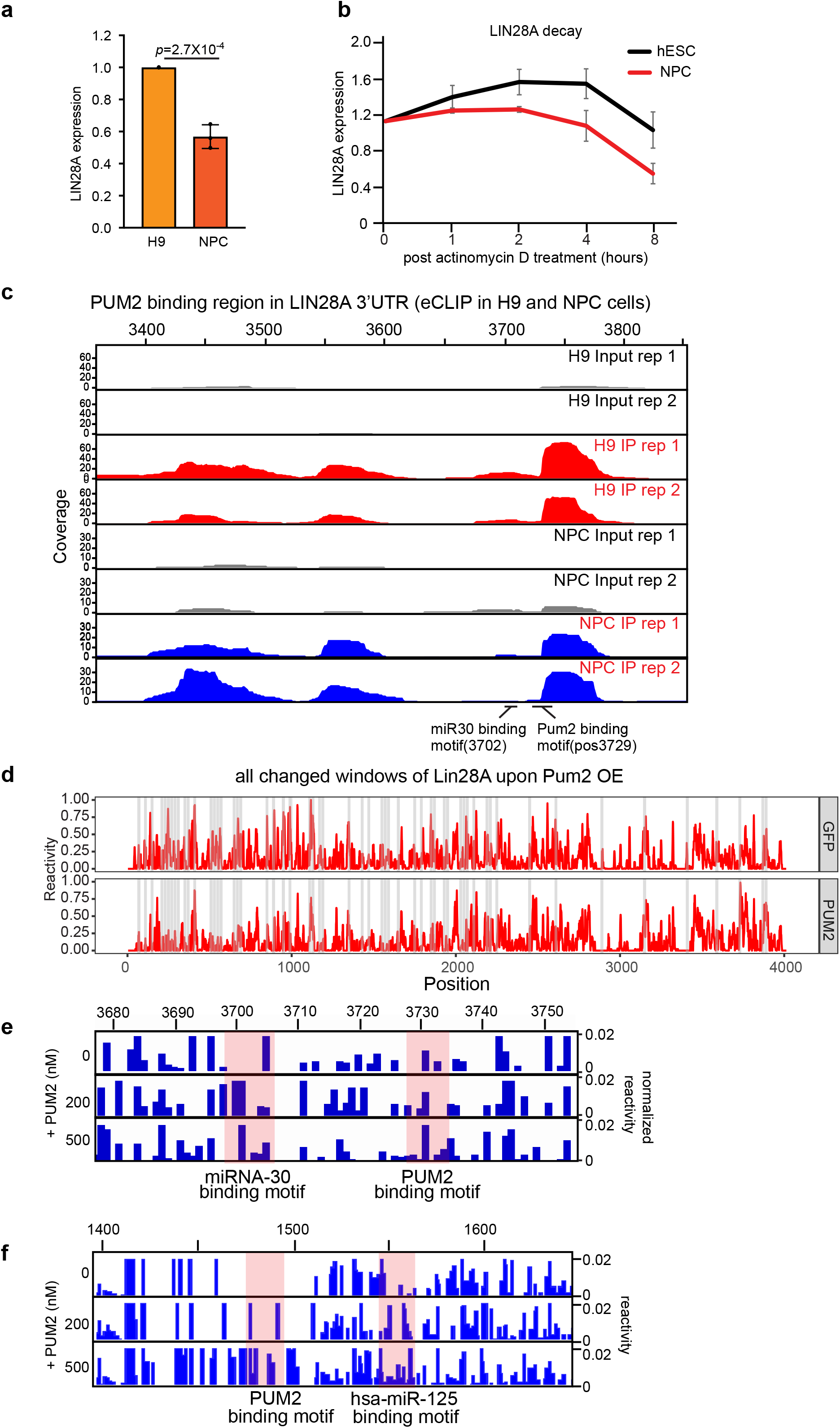
PUM2 and miR-30 regulated LIN28A through structure. **a,** Barplot showing the relative expression level of LIN28A in hESC (H9, yellow) and NPC (orange). **b,** Gene expression of LIN28A upon 0, 1, 2, 4, 8 hours post actinomycin D treatment in hESC (black line) and in NPC (red line). **c,** the depth of coverages obtained from eCLIP experiments of PUM2 in hESC and NPC cells on the 3’UTR region of gene LIN28A (from pos 3400 to 3800) were shown. The PUM2 binding site (position 3729) and miR-30 binding site (position 3702) are labeled. **d,** icSHAPE reactivity profile of LIN28A upon overexpression of GFP (top) or PUM2 (bottom). The grey bars are the reactivity changing regions between GFP and PUM2 overexpression. **e, f,** in vitro SHAPE-MaP reactivity profiles of PUM2 and miR-30 (**e**) or miR-125 (**f**) binding regions along LIN28A in the presence of increasing concentrations of PUM2 (0, 200nM, 500nM).

**Supplementary Figure 15.**
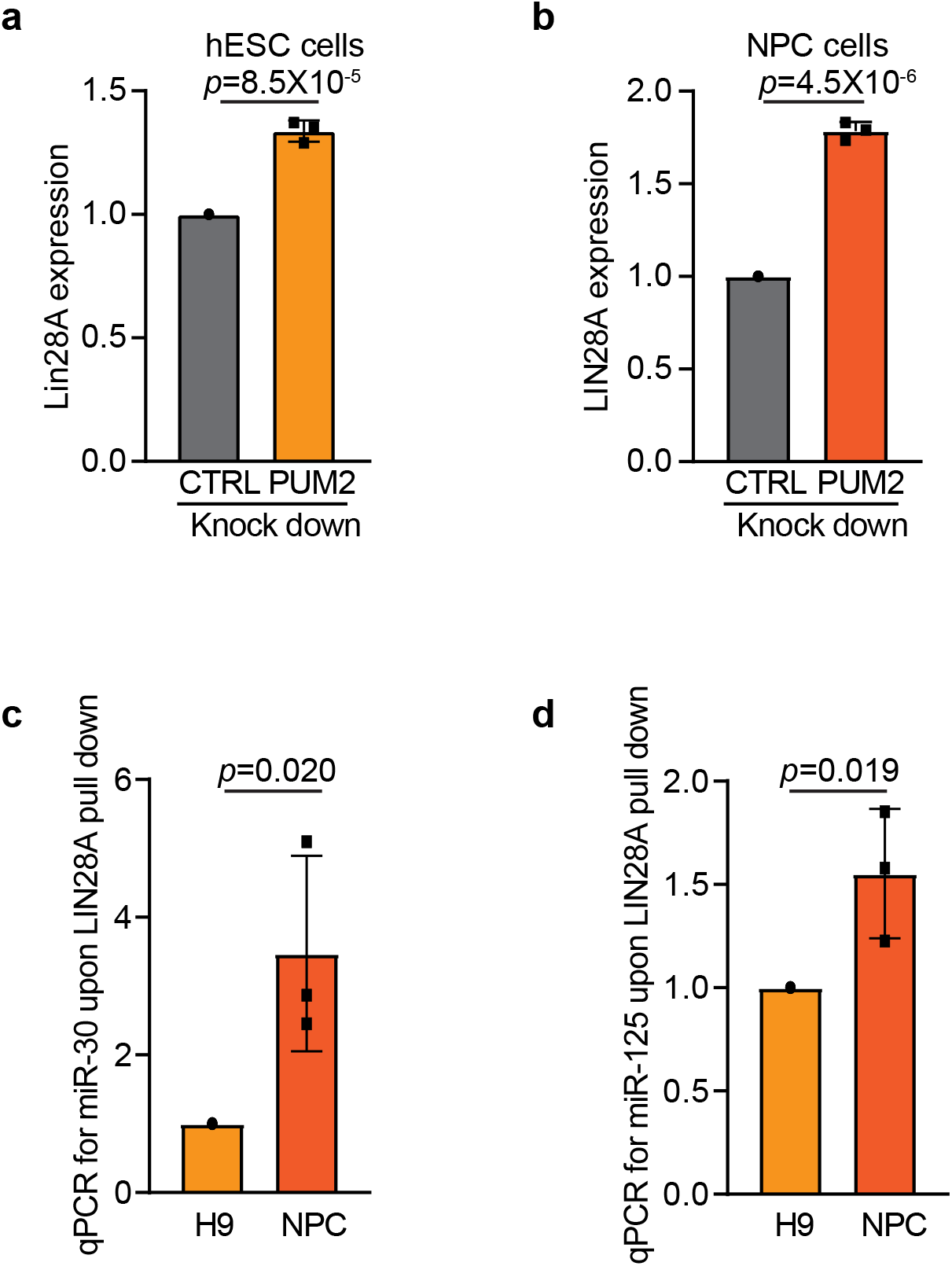
Impact of PUM2 binding on LIN28A. a, b, Barplots showing the expression levels of LIN28A after PUM2 knock down in hESC cells (**a**) and NPC cells (**b**). **c, d,** Barplots showing the amount of miR-30 (**c**) and miRNA-125 (**d**) associated with LIN28A upon pulldown of LIN28A, using biotinylated probes, in hESC and NPC cells.

**Supplementary Figure 16.**
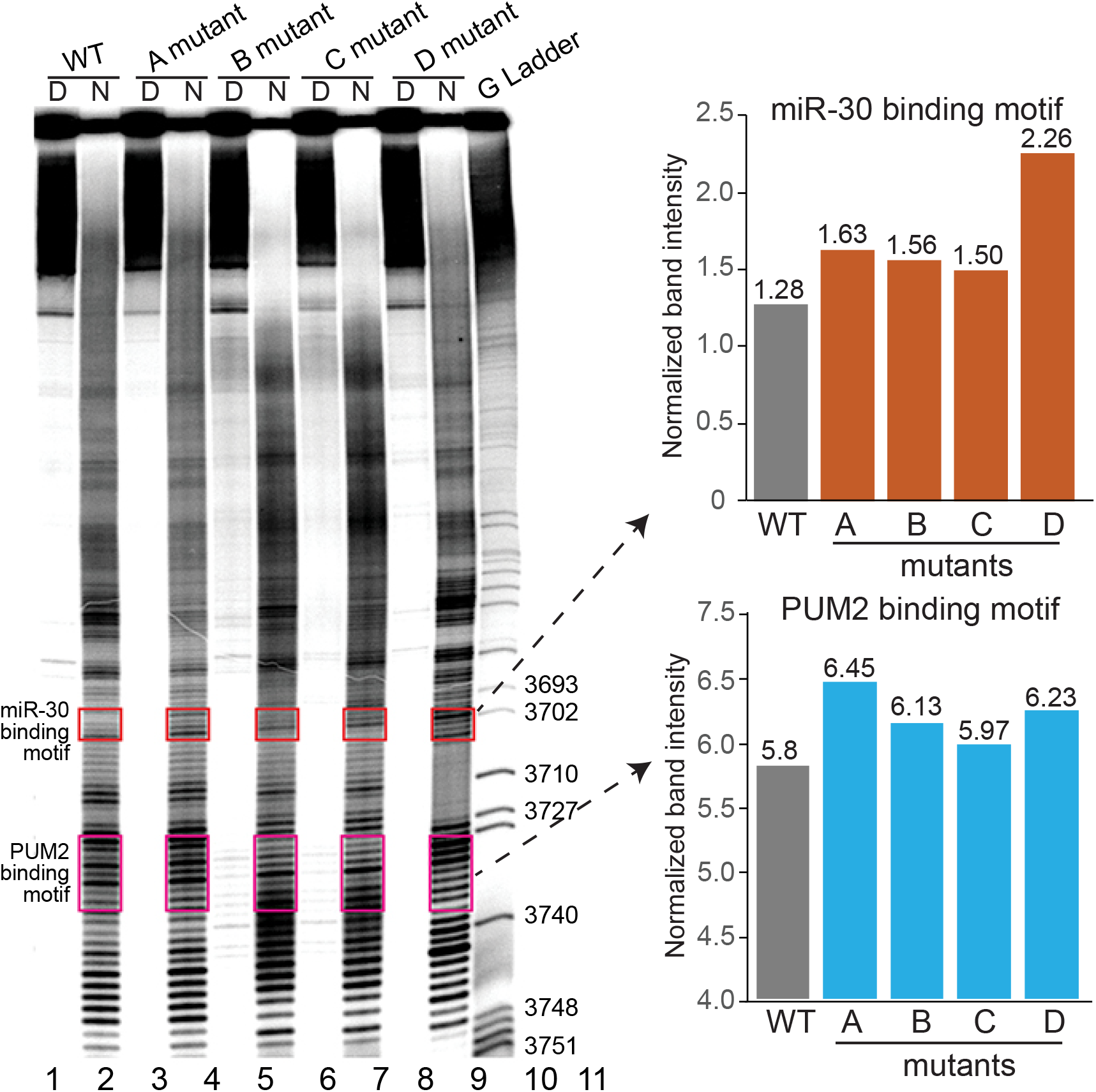
Foot printing of mutated RNA sequences. Sequencing gel showing RT stoppages from WT and mutant DMSO and NAI-N3 treated RNAs. The G ladder is in lane 11, WT DMSO and NAI-N3 treated samples are in lanes 1 and 2 respectively, Mutant A DMSO and NAI treated samples are in lanes 3 and 4, Mutant B DMSO and NAI treated samples are in lanes 5 and 6, compensatory Mutant C DMSO and NAI treated samples are in lanes 7 and 8, and Mutant D DMSO and NAI treated samples are in lanes 9 and 10. miR-30 binding motif and PUM2 binding motif region were boxed in red and pink respectively. The band intensity of the boxed images were measured using ImageJ. The intensity of the miR-30 RNA binding site in WT and mutants are shown as barcharts on the top right. The intensity of the PUM2 RNA binding site in WT and mutants are shown as barcharts on the bottom right.

